# RNA Profiling of Neuropathic Pain-Associated Human DRGs Reveal Sex-differences in Neuro-immune Interactions Promoting Pain

**DOI:** 10.1101/2021.11.27.470190

**Authors:** Pradipta R. Ray, Stephanie Shiers, Diana Tavares-Ferreira, Ishwarya Sankaranarayanan, Megan L. Uhelski, Yan Li, Robert Y. North, Claudio Tatsui, Gregory Dussor, Michael D. Burton, Patrick M. Dougherty, Theodore J. Price

## Abstract

Neuropathic pain is a leading cause of high impact pain, is often disabling and is poorly managed by current therapeutics. Here we focused on a unique group of neuropathic pain patients undergoing thoracic vertebrectomy where the DRG is removed as part of the surgery allowing for molecular characterization and identification of mechanistic drivers of neuropathic pain independently of preclinical models. Our goal was to quantify whole transcriptome RNA abundances using RNA-seq in pain-associated human DRGs from these patients, allowing comprehensive identification of molecular changes in these samples by contrasting them with non-pain associated DRGs. We sequenced 70 human DRGs, including over 50 having mRNA libraries with neuronal mRNA. Our expression analysis revealed profound sex differences in differentially expressed genes including increase of *IL1B*, *TNF*, *CXCL14*, and *OSM* in male and including *CCL1*, *CCL21*, *PENK* and *TLR3* in female DRGs associated with neuropathic pain. Co-expression modules revealed enrichment in members of JUN-FOS signaling in males, and centromere protein coding genes in females. Neuro-immune signaling pathways revealed distinct cytokine signaling pathways associated with neuropathic pain in males (OSM, LIF, SOCS1) and females (CCL1, CCL19, CCL21). We validated cellular expression profiles of a subset of these findings using RNAscope *in situ* hybridization. Our findings give direct support for sex differences in underlying mechanisms of neuropathic pain in patient populations.

## Introduction

Neuropathic pain affects millions of US adults, is a primary cause of high impact chronic pain and is poorly treated by available therapeutics (Finnerup et al., 2015; Dahlhamer et al., 2018; Pitcher et al., 2018). Pre-clinical studies in rodents have identified important roles of neuronal plasticity (Price and Gold, 2018; Price and Ray, 2019) and neuro-immune interactions (Ji et al., 2016; Sommer et al., 2018) in neuropathic pain, but molecular and anatomical differences between rodent models and humans (Davidson et al., 2016; Ray et al., 2018; Rostock et al., 2018; Shiers et al., 2020; Middleton et al., 2021; Shiers et al., 2021; Tavares-Ferreira et al., 2021) suggest that the (diverse) biological processes involved in the chronification and maintenance of human pain remain ill-understood, and therapeutics based on rodent models have faced serious translational challenges (Price et al., 2018; Renthal et al., 2021). Molecular mechanisms of neuropathic pain in patients need to be understood and putative drug targets identified in order to develop the therapeutics that can meet this medical challenge.

Here we have built upon our work with patients undergoing thoracic vertebrectomy surgery which often involves removal of dorsal root ganglia (DRGs) (North et al., 2019). This provides an opportunity to identify neuropathic pain in specific dermatomes prior to surgery allowing comparison of DRGs associated with neuropathic pain to those without. Our goal was to quantify whole transcriptome RNA abundances using RNA-seq in pain-associated DRGs to comprehensively identify differences in RNA profiles linked to the presence of neuropathic pain in male and female patients. Our previous work demonstrated transcriptomic differences in human DRG (hDRG) associated with neuropathic pain, but our sample size was insufficient to reach direct conclusions about sex differences in underlying neuropathic pain mechanisms (North et al., 2019). Given the overwhelming evidence for such differences in preclinical neuropathic pain models (Sorge et al., 2015; Rosen et al., 2017; Inyang et al., 2019; Mogil, 2020; Yu et al., 2020; Agalave et al., 2021), we hypothesized that an increased sample size would give us the ability to detect clear differential expression of neuro-immune drivers of neuropathic pain in this patient population.

In this study, we have sequenced 70 hDRGs from 40 thoracic vertebrectomy patients, with 51 having mRNA libraries that are not de-enriched in sensory neuronal mRNA. The dataset is the largest repository of hDRG RNA profiles described to date (dbGaP id will be added upon publication) and has allowed us to perform a sex-stratified transcriptome-wide association study (TWAS), followed by co-expression module analysis of neuropathic pain-associated genes. We additionally performed integrative analysis with existing hDRG gene expression, protein interaction and protein functional annotation databases for identifying putative cell type of expression and function of these genes. Our analysis revealed clear sex differences in sets of genes associated with neuropathic pain in men and women. These gene products are prominently involved in neuro-immune signaling and neuronal plasticity. We validated the cell type expression of a subset of these genes with RNAscope *in situ* hybridization. Our findings paint a picture of neuro-immune signaling that likely drives neuropathic pain in this patient population.

## Materials and Methods

### Consent, tissue and patient data collection

All protocols were reviewed and approved by the UT Dallas (UTD) and MD Anderson Cancer Center (MDACC) Institutional Review Boards. All protocols and experiments conform to relevant guidelines and regulations, in agreement with the Declaration of Helsinki. Patients undergoing thoracic vertebrectomy at MDACC for malignant tumors involving the spine were recruited as part of the study. Informed consent for participation was obtained for each patient during study enrolment.

All donors were undergoing surgery which required ligation of spinal nerve roots for spinal reconstruction or tumor resection. Tissue extraction and patient data collection was performed as described in North *et al* (North et al., 2019). In short, spinal roots were cut, the ganglia immediately transferred to cold (∼4C) and sterile balanced salt solution containing nutrients, taken to the laboratory, cleaned and partitioned into three or four sections (typically quartered). One section was frozen in RNAlater and shipped on dry ice to UTD for RNA sequencing (RNA-seq).

Data (including demographics, clinical symptoms and medical history) were obtained from consented patients at MDACC through retrospective review of medical records and prospective data collection during study enrolment. Neuropathic pain was defined as a binary clinical variable for purposes of reporting consistency. The presence or absence of neuropathic or radicular pain for each dermatome was performed in a manner consistent with the guidelines for (definite or probable) neuropathic pain from Assessment Committee of the Neuropathic Pain Special Interest Group of International Association for the Study of Pain (IASP) (Haanpää et al., 2011; Jensen et al., 2011). A harvested hDRG was determined to be associated with neuropathic or radicular pain if the patient had documented spontaneous pain, hyperalgesia, or allodynia in a region at or within two classically defined dermatomes of the harvested ganglion in question and was considered not to be associated with neuropathic pain if not (or if the harvested ganglion was from the side contralateral to reported pain in a patient with only unilateral symptoms). Remaining scenarios were categorized as inconclusive. All pain reports dated back at least one month, with one exception (66T12R). Most subjects were cancer patients treated with chemotherapeutics but the DRGs collected and analyzed did not have any signs of tumor and only a few were associated with dermatomes affected by length-dependent neuropathy. De-identified patient data, including usage of drug treatment history, for the entire cohort can be found in **Table 1** and **Supplemental File 1**.

**Table 1.**
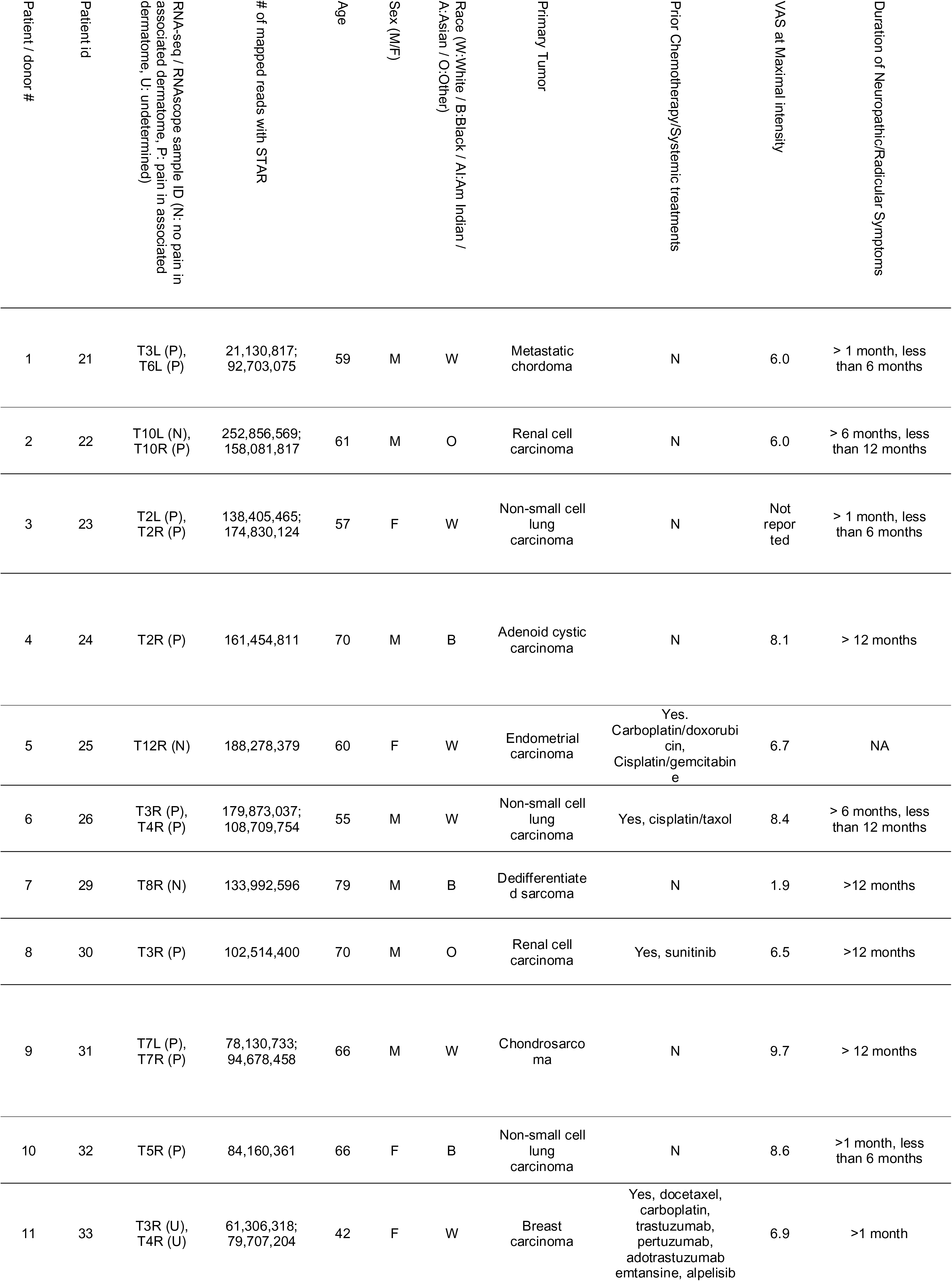
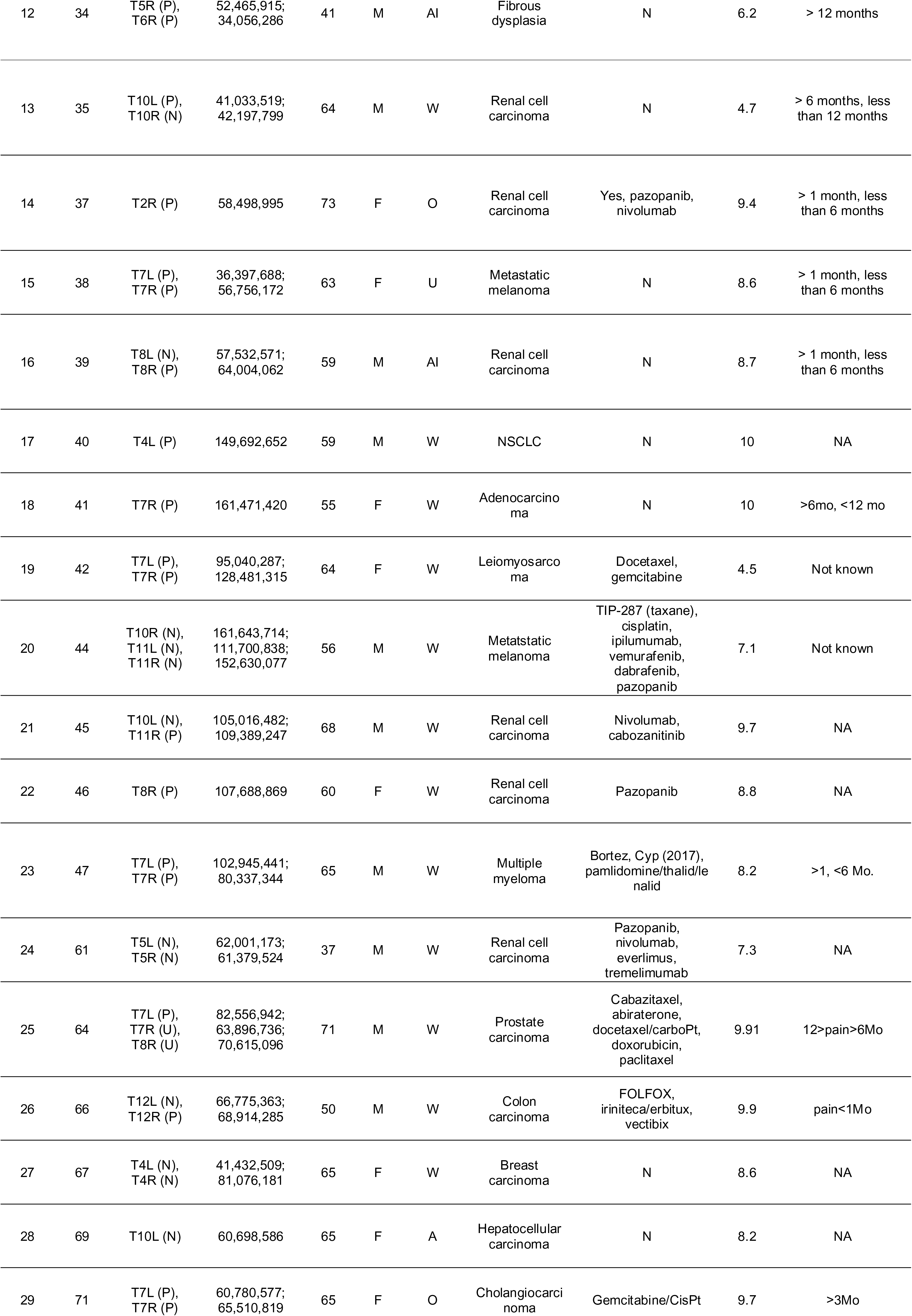
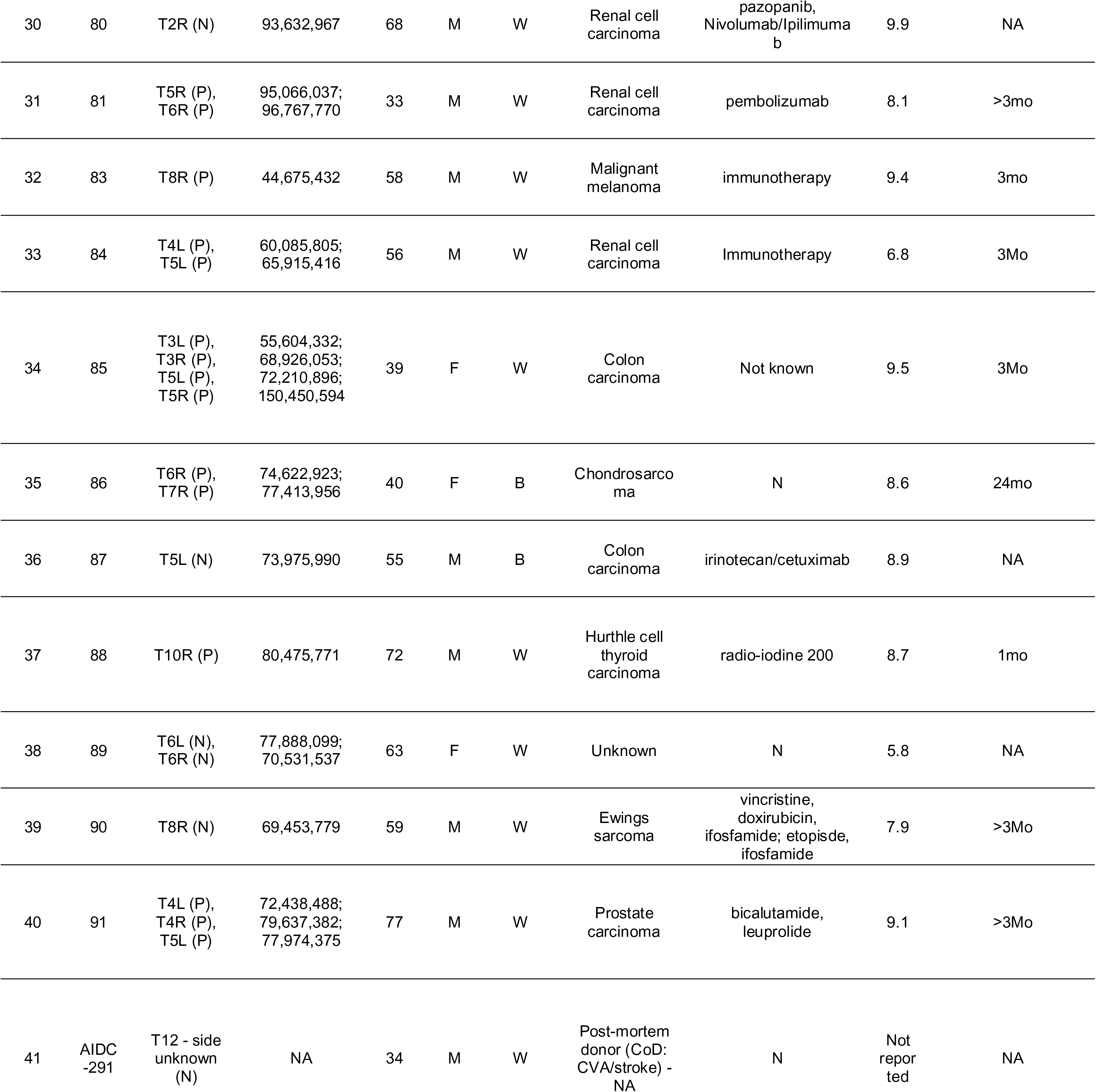
Patient details table. Relevant demographic, clinical and medical history variables of de-identified patients

### RNA-seq library preparation, mapping and abundance quantification

Total RNA from each DRG sample were purified using TRIzol^TM^ and depleted of ribosomal RNA. RNA integrity was assessed and Illumina Tru-seq library preparation protocol was used to generate cDNA libraries according to manufacturer’s instructions. Single-end sequencing of each library was performed in multiplexed fashion across several batches as samples became available on the Illumina Hi-Seq sequencing platform. Sequenced reads were trimmed to avoid compositional bias and lower sequencing quality at either end and to ensure all quantified libraries were mapped with the same read length (38bp), and mapped to the GENCODE reference transcriptome (v27) (Frankish et al., 2019) in a strand-aware and splicing-aware fashion using the STAR alignment tool (Dobin et al., 2013). Stringtie (Pertea et al., 2015) was used to generate relative abundances in Transcripts per Million (TPM), and non-mitochondrial coding gene abundances were extracted and re-normalized to a million to generate coding TPMs for each sample (**Supplemental File 2, Sheet A**).

In our previous study (North et al., 2019), we noted variation in enrichment of neuronal mRNA content per sample, likely caused primarily by technical factors (what proportion of the neuronal cell bodies and axon, as opposed to myelin, perineurium, and epineurium are sampled in each quartered DRG, and the amount of viable mRNA extracted from these). With the number of samples increasing almost four-fold in our present study, we identified a greater spread in neuronal mRNA content across samples. Irrespective of whether such variation is due to biological or technical causes, transcriptome-wide association studies (TWAS) would be confounded by de-enrichment of neuronal mRNA in a subset of samples (that would likely contribute to both within-group and between-group variation, when grouped by pain state). Thus, samples that show moderate or strong neuronal mRNA de-enrichment were excluded from downstream analysis.

Based on a panel of 32 genes that are known neuronal markers (like *RBFOX3*) or DRG-enriched in human gene expression (Ray et al., 2018), and further validated to be enriched in neuron-proximal barcodes in Visium spatial sequencing for hDRG (Tavares-Ferreira et al., 2021), we tabulated the relative abundance (in Transcripts per Million or TPMs) of these genes in each of our samples (**Supplemental File 2, Sheet B**). Quantiles were calculated across samples for each gene, and a subset of the samples showed low quantiles for the vast majority of the genes in the gene panel, suggesting that the low gene expression was not the result of down-regulation, but systematic de-enrichment of neuronal mRNAs. The neuronal mRNA enrichment index for each sample was calculated as a median of the per-gene quantile value (median calculated across genes in the gene panel). Based on the neuronal mRNA enrichment index (ranges for the index for each group shown in **Supplemental File 2, Sheet B**), we grouped the samples into two groups: samples with moderately or strongly de-enriched neuronal mRNA, and samples with a higher proportion of neuronal mRNA. Out of 70 samples, 51 RNA abundance profiles with higher neuronal mRNA content were retained for downstream analysis. We also found that the neuronal mRNA enrichment index was correlated with the expression profile of 1239 genes (Pearson’s R > 0.55, uncorrected *p* < 0.00005), many of which are known to be neuronally enriched in the hDRG (**Supplemental File 2, Sheet C**). Gene TPMs for these 51 samples are presented in **Supplemental File 2, Sheet D**. Of these, the dermatome of one (64T8R) sample could not be conclusively placed into pain or no pain categories, so this sample was also not used for downstream analysis. All of our samples used for downstream analysis, had an adequate (> 40 million) number of uniquely mapped reads, as shown in **Table 1**, and have adequate library complexity (> 12,000 genes detected, **Supplemental File 2, Sheet D**).

Finally, in order to standardize TPM distributions across samples, we performed two quantile normalizations – for samples with > 15,500 genes detected, and for samples with 13,400 to 14,100 genes detected (25T12R and 30T3R). We reset the abundances for genes with zero counts to zero after quantile normalization. Quantile normalization caused the TPM distribution across samples to be similar to each other, reducing variance from technical factors like sequencing depth, especially for higher quantiles (0.33 or higher). Quantile normalized TPMs are presented in **Supplemental File 2, Sheet E**.

### Transcriptome-wide association study with neuropathic pain

We stratified the sample set by sex. Our analysis identifies sample-level TWAS between the digital readouts of relative abundances of coding genes, with the binary variable representing. Dermatome-associated pain. In experiments on rodent models with identical genetic landscapes, and consistent insult or injury models defining groups, a straightforward way to perform TWAS is to directly perform differential expression analysis (with sex as a batch or partitioning variable) (Anders and Huber, 2010). Such a straightforward approach is unlikely to work in real world human transcriptome datasets for a variety of reasons. Gene expression of pain-associated genes are likely to occupy a spectrum, due to the fact that there are some differences in pathologies, and cell type proportions from sample to sample, differences in genetic landscape, and variations in medical and clinical history from patient to patient. The heterogeneous nature of neuropathic pain suggests multiple molecular mechanisms, some of which may be post-transcriptional and thus not be detectable by sampling steady-state transcriptomes. Further, a few of the DRGs not associated with pain could also show pain-like molecular signatures due to the fact that proximal mammalian dermatomes can overlap (Rigaud et al., 2008).

Thus, instead of a traditional differential expression (DE) analysis, we identify distributional shifts in abundance between pain and non-pain samples, in a sex-stratified fashion. For each of the sexes, we contrasted gene expression between the pain and non-pain sub-cohorts. Such an approach accounts not just for changes in the mean or median, but also for changes in variance in the disease state (Ho et al., 2008), a potential confounding effect in such studies.

Since our libraries were constructed from total RNA, we removed from downstream analysis genes that did not have a validated peptide sequence, fusion genes, or families that have large numbers of pseudogenes like olfactory receptors (Menashe et al., 2006) even though they may be predicted as coding genes in the reference annotation. To ensure consistently detectable genes with lower sampling variance, we constrained our analysis to genes with TPMs > 0.5 qnTPM (quantile normalized TPM) at the median (50^th^ percentile), and > 1.0 qnTPM at the upper quartile (75^th^ percentile) for the sub-cohort that was tested for increased gene expression (rounded to one decimal place).

In our previous experience in analyzing sex-differential RNA-seq expression in large human cohorts (Ray et al., 2019), we found the most distinct differences occurred between the median and upper quartile (75^th^ percentile). Here, we identified distributional shifts in gene abundance between the pain and non-pain sub-cohorts in each sex by comparing order statistics between the 20^th^ and 80^th^ percentiles (to partially mitigate the effects of outliers). We constrained that the directionality of change in abundance (increase or decrease in pain vs non-pain sub-cohorts) had to be consistent between the median, upper quartile, and the maximum analyzed quantile (80^th^ percentile) – which might suggest consistency with a phenotype or degree of pathology. We then selected for genes with the largest changes in abundance between pain states by filtering out genes with median fold change < 1.5, or with a maximum fold change in the top two quartiles < 2.0 (up to a rounding error in the fold change of one decimal point). A smoothing factor of 0.1 was added to both the numerator and denominator to calculate fold changes. Finally, to identify genes with maximal distributional shift, we filtered based on two metrics. First, to identify genes with the largest difference in values for the same quantiles, we constrained the gene set to genes with area between the quantiles of the distributions (between the pain and non-pain sub-cohorts of one sex) > 5% of the total area of the quantile plot. The approximate normalized signed area between the quantile curves was calculated as follows:

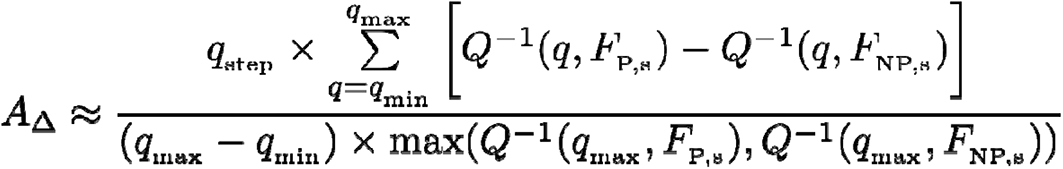

where Q^-1^ is the inverse quantile function, F_P,s_ and F_NP,s_ are the empirically estimated distributions for the pain and non-pain sub-cohort respectively for the sex s in question, and q changes from q_min_ (0.2) to q_max_ (0.8) in steps of 0.025 (q_step_). Then, to identify genes with the largest difference in quantiles for the same values, we calculated the shift in quantiles for each of the three quartiles in the distribution of a sub-cohort and summed them, and filtered out genes with a negative sum:

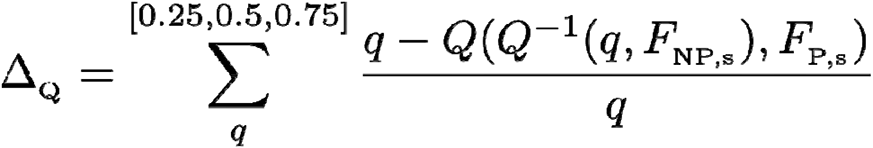

where Q is the quantile function, Q^-1^ is the inverse quantile function, F_P,s_ and F_NP,s_ are the empirically estimated distributions for the pain and non-pain sub-cohort respectively for the sex s in question respectively. For the filtering criteria in genes that are increased in non-pain cohorts with respect to pain cohorts for males or females, F_P,s_ and F_NP,s_ are interchanged in the formulae. The genes that were increased in the pain and non-pain cohorts for each sex are presented in **Supplemental File 3, Sheets A – D**. We refer to the genes that are increased or decreased in abundance in the pain cohorts as pain-associated genes.

Additionally, for four male patients with unilateral pain (22, 35, 39 and 66), we have ipsilateral and contralateral DRG samples. These samples lend themselves to a classical paired DE analysis with paired fold changes, since medical and clinical history, as well as genetic background is consistent across each pair and were found to be consistent in terms of the genes upregulated in the pain samples. Genes whose median log fold change across the four pairs were two-fold or greater, and are presented in **Supplemental File 2, Sheets E and F**. We limited our analysis to highly expressed genes - more than 2 (out of 4) of the samples in the group being tested for increased gene expression were required to have qnTPM > 2.0.

### Co-expression module analysis

In order to identify transcriptional programs that may drive changes in gene abundance or identity of cell types where these pain-associated genes were expressed, we identified co-expression modules (based on quantile-normalized TPMs) for genes that were increased in pain or no-pain cohorts, for each sex. Pearson’s R was calculated for each pain-associated gene in the relevant sex, and genes with a statistically significant Pearson’s R that were highly expressed (median qnTPM > 2.0) in the sex-stratified sub-cohort where the co-expression module was increased in abundance, were retained as members of the module. There were several co-expression modules with > 100 genes, but in our study, we focused on two large co-expression modules (gene co-expressing with *OSM* in males, and with *IFIT1* in females) that showed statistically significant Pearson’s R, and enrichment for well-known signaling pathways. Further, transcriptional regulators that are part of these modules were also profiled in order to better understand the transcriptional identity of the cells where these genes were expressed, and potential regulatory cascades that may be implicated. **Supplementary Figures 1A and B**, and **Supplemental File 3, Sheets G and H** show transcription factor genes (R > 0.55) and all genes (R > 0.76) that were co-expressed with *OSM*, respectively. **Supplementary Figures 2A and B**, and **Supplemental File 3, Sheets I and J** show transcription factor genes (R > 0.55) and all genes (R > 0.76) that were co-expressed with *IFIT1*, respectively.

The threshold for Pearson’s R chosen in the male and female cohorts were as follows: R > 0.76 for whole transcriptome correlation corresponds to *p* < 7.1e-7 in males [N=31], and to *p* < 1.6e-4 in females (N=19) for a two-tailed test). We chose a less stringent threshold for identifying co-expressed transcription factor (TF) genes, since < 1000 transcription factors are expressed in most cell types (as opposed to ∼ 10,000 genes for the whole transcriptome). R > 0.55 for TF co-expression analysis corresponds to *p* < 6.7e-4 in males (N=31), and to *p* < 7.3e-3 in females (N=19) for a two-tailed test. For each gene in the co-expression module, 80% of the samples were randomly sampled repeatedly (N = 20), and small (0.01) values were randomly added or subtracted to each element of the vector to ensure that randomly detected low expression or outlier samples were not driving the correlation.

### Functional annotation and overlap analysis

The list of transcription factors was obtained from literature (Lambert et al., 2018) and signaling pathway gene sets were obtained from the MSigDB Hallmark database (Liberzon et al., 2015). Gene set enrichment analysis was performed with Enrichr (Kuleshov et al., 2016). We identified potential protein interaction using STRINGdb (Jeanquartier et al., 2015). We used products of pain-associated genes that were increased in pain, genes of the profiled co-expression modules, and hDRG-expressed genes (Ray et al., 2018) that are known to interact with them to seed the network, using only linkages with medium or higher confidence, for gene product pairs with known molecular interaction or co-expression.

### RNAscope assay

Based on our findings from the RNA-seq analysis, we decided to validate the gene expression of identified pain-associated genes and identify their cell types of expression using RNAscope *in situ* hybridization assay. We chose two pain-associated samples and one non-pain sample for each sex for performing RNAscope. Since RNA-seq is a destructive method, RNAscope was performed on a different piece of the same harvested DRG sample (from which the RNA-seq was conducted) that had been previously banked.

Given the limited availability of banked fresh-frozen hDRG sections, we used the following tissues in our RNAscope experiments: Male 84T4L (pain; thoracic 4 left), male 91T5L (pain; thoracic 5 left), female 85T3L (pain; thoracic 3 left), female 86T6R (pain; thoracic 6 right), female 89T6R (non-pain; thoracic 6 right). We had no banked fresh frozen non-pain thoracic hDRG samples collected from vertebrectomy patients, so we used a thoracic-12 hDRG from a non-pain organ donor (AIDC291) acquired through our collaboration with the Southwest Transplant Alliance, recovered from an organ donor with no history of neuropathic pain (**Table 1**). Thus, this sample does not have a corresponding RNA-seq assay.

RNAscope *in situ* hybridization multiplex version 2 on fresh frozen tissue was performed as instructed by Advanced Cell Diagnostics (ACD). A 2-minute protease IV digestion was used for all experiments and fluorescin, Cy3 and Cy5 dyes (Akoya) were used in lieu of Opal dyes. The probes used were: *OSM* (ACD Cat 456381), *IL1B* (ACD Cat 310361), *TNF* (ACD Cat 310421), *IL32* (ACD Cat 541431-C2), *IFIT1* (ACD Cat 415551), *AIF1* (ACD Cat 433121-C3), *HLA-DQB1-O1* (ACD Cat 527021-C2), *TRPV1* (ACD Cat 415381-C1, C2). All tissues were checked for RNA quality by using a positive control probe cocktail (ACD Cat 320861) which contains probes for high, medium and low-expressing mRNAs that are present in all cells (ubiquitin C > Peptidyl-prolyl cis-trans isomerase B > DNA-directed RNA polymerase II subunit RPB1). All tissues showed robust signal for all 3 positive control probes. A negative control probe (ACD Cat 320871) against the bacterial *DapB* gene (ACD) was used to check for non-specific/background label. 40X images were acquired on an FV3000 confocal microscope (Olympus). The acquisition parameters were set based on guidelines for the FV3000 provided by Olympus.

## Results

We analyzed neuronal enrichment of 70 DRGs collected from thoracic vertebrectomy patients (patient information shown in **Table 1** and **Supplemental File 1**). We tabulated relative abundance of all non-mitochondrial coding genes in the hDRG for each sample (**Supplemental File 2, Sheet A**) and based on the abundance of neuronally enriched genes, calculated the neuronal mRNA enrichment index for each sample (**Supplemental File 1, Sheet B**). We grouped the samples in two: samples with moderately or strongly de-enriched neuronal mRNA, and samples with a higher proportion of neuronal mRNA (**Supplemental File 2, Sheet B)**. 1239 genes we found to be strongly correlated to the neuronal mRNA enrichment index, including most known sensory neuron enriched genes (including *P2RX3, SCN9A, SCN10A, SCN11A, CALCA* and *CALCB*) (**Supplemental File 2, Sheet C**). We identified 51 samples with higher proportions of neuronal mRNA and these RNA abundance profiles were retained for downstream analysis (**Supplemental File 2, Sheet D).** All samples had sufficient library complexity (> 12,000 coding genes detected). Quantile-normalization was performed on all samples to account for sample-to-sample differences in library complexity and sequencing depth (qnTPMs for retained samples provided in **Supplemental File 2, Sheet E**).

### Transcriptome-wide association of gene expression with neuropathic pain

Based on our association analysis of gene abundances, we found that 195 genes were increased (pain-associated) and 70 genes were decreased in male DRG samples with dermatomes associated with neuropathic pain (**Supplemental File 3, Sheets A and B**), while 576 genes were increased (pain-associated) and 254 genes decreased in pain-associated female DRG samples (**Supplemental File 3, Sheets C and D**). The higher number of female pain-associated genes is likely due to using consistent thresholds for effect sizes across sexes, while the ratio of the number of samples to the number of subjects in the female pain sub-cohort (12:9 = 1.33) is higher than the corresponding ratio in the male pain sub-cohort (19:15 = 1.26), leading to lower within-group variability.

Both male (including *IL1B*, *TNF*, *CXCL14*, *OSM, EGRB, TRPV4, LIF, CCL3/4*) and female (including *CCL1, CCL19, CCL21*, *PENK, TRPA1, ADORA2B, GLRA3*) pain-associated genes showed a strong enrichment of genes that are well known to be pain, nociception or inflammation related (LaCroix-Fralish et al., 2011). The top pain-associated genes (ranked by the area between the quantile curves of each pain state) that are increased in the pain state in males and females are shown in Figures 1 and 2, respectively. Strikingly, at the same effect size, only a handful of the male and female genes with pain-associated increases overlapped, including *ACAN, CPXM1, DIO2, FNDC1, HAMP, IQGAP3, LAIR2, LILRA5, LMNB1, LRRC24, LY6G5C, SIGLEC7* and *TGM*, with only *HAMP* among the top 25 pain-associated genes for either sex to have a median fold change of > 2.0 for both sexes (Figure 3A). This suggests that the enriched immune cell types and regulatory pathways in these sex-stratified sub-cohorts are strikingly different and raises the possibility that neuronal changes driven by neuro-immune interaction may be sex differential too.

**Figure 1.**
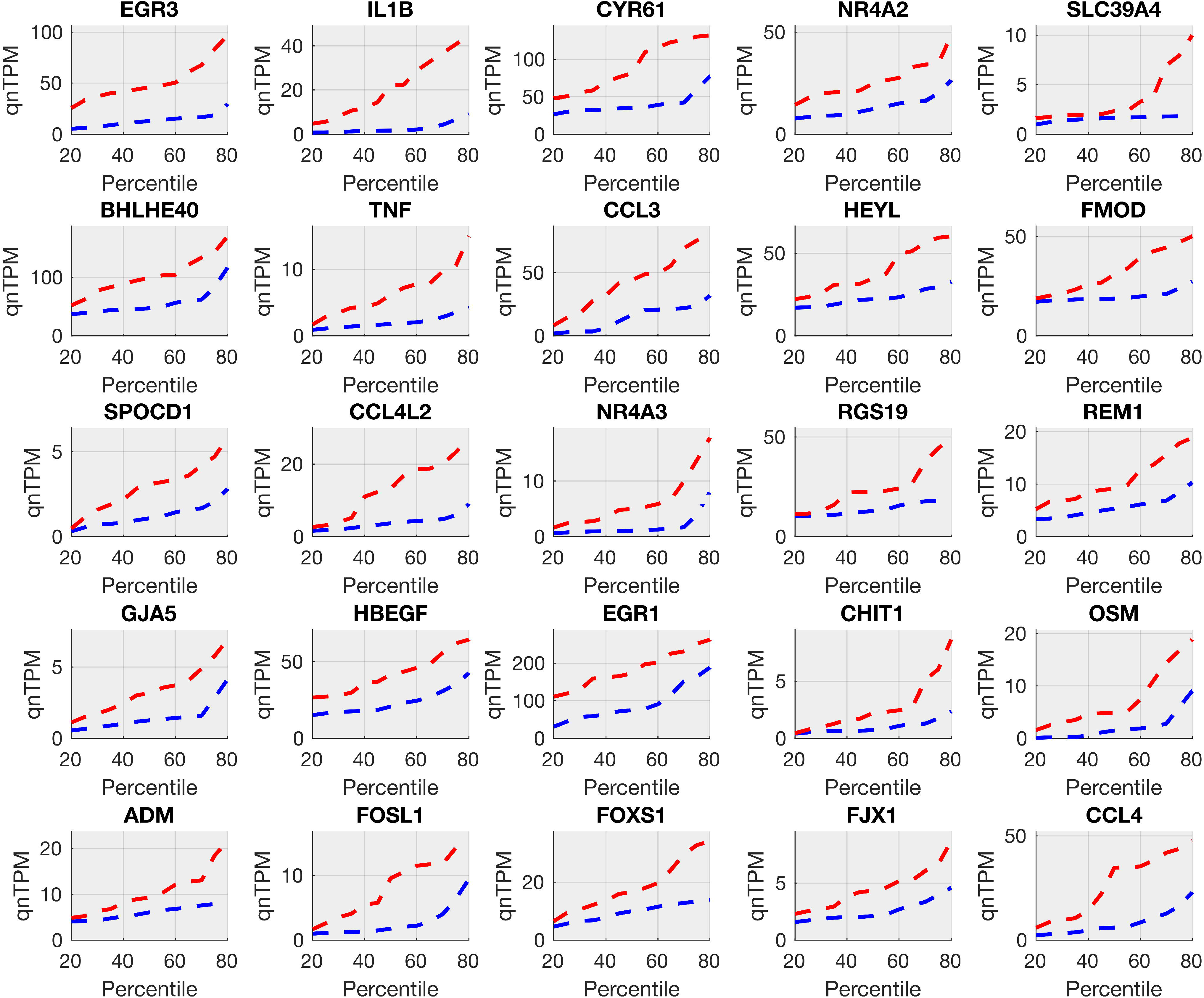
Top 25 pain-associated genes in the male cohort (increased in pain). Pain-associated genes in male samples (**Supplemental File 3, Sheet A**), show systematic increases in pain for gene abundance quantiles of multiple members of AP-1 signaling (*EGR3, FOSL1*), pro-inflammatory cytokines (*IL1B, CCL3, CCL4*), TNF signaling (*TNF, IL1B*) and other transcriptional regulators (*NR4A2, FOXS1, HBEGF*) relevant to the peripheral nervous system and pain.

**Figure 2.**
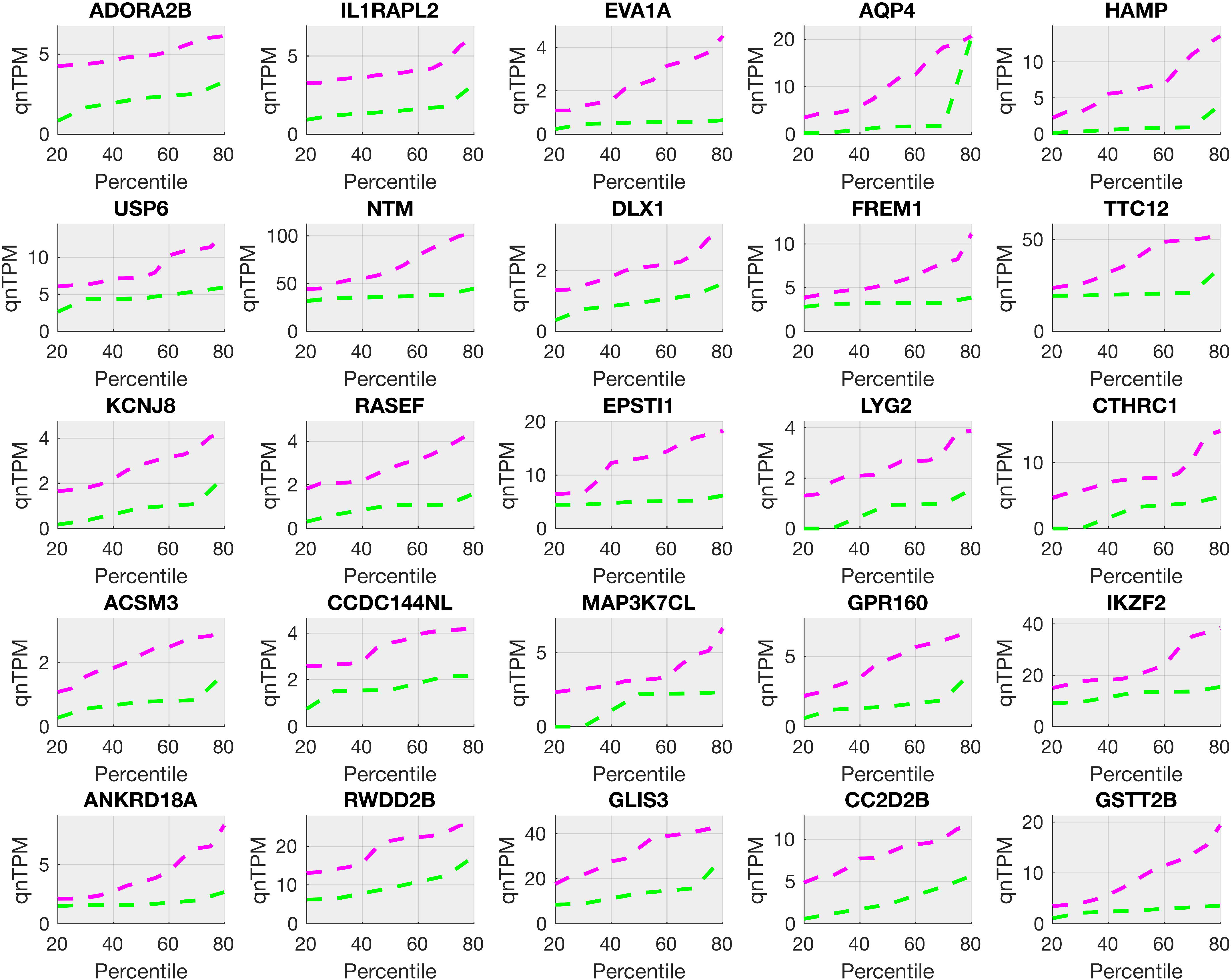
Top 25 pain-associated genes in the female cohort (increased in pain). Pain-associated genes in female samples (**Supplemental File 3, Sheet C**), show systematic increases in pain for gene abundance quantiles of multiple members of receptor genes (*ADORA2B, IL1RAPL2, GPR160*), pro-inflammatory and proliferation-related genes (*HAMP, FREM1*), vesicular trafficking genes (*LYG2, RASEF*), and interferon-response genes (*USP6, TTC12*)

**Figure 3.**
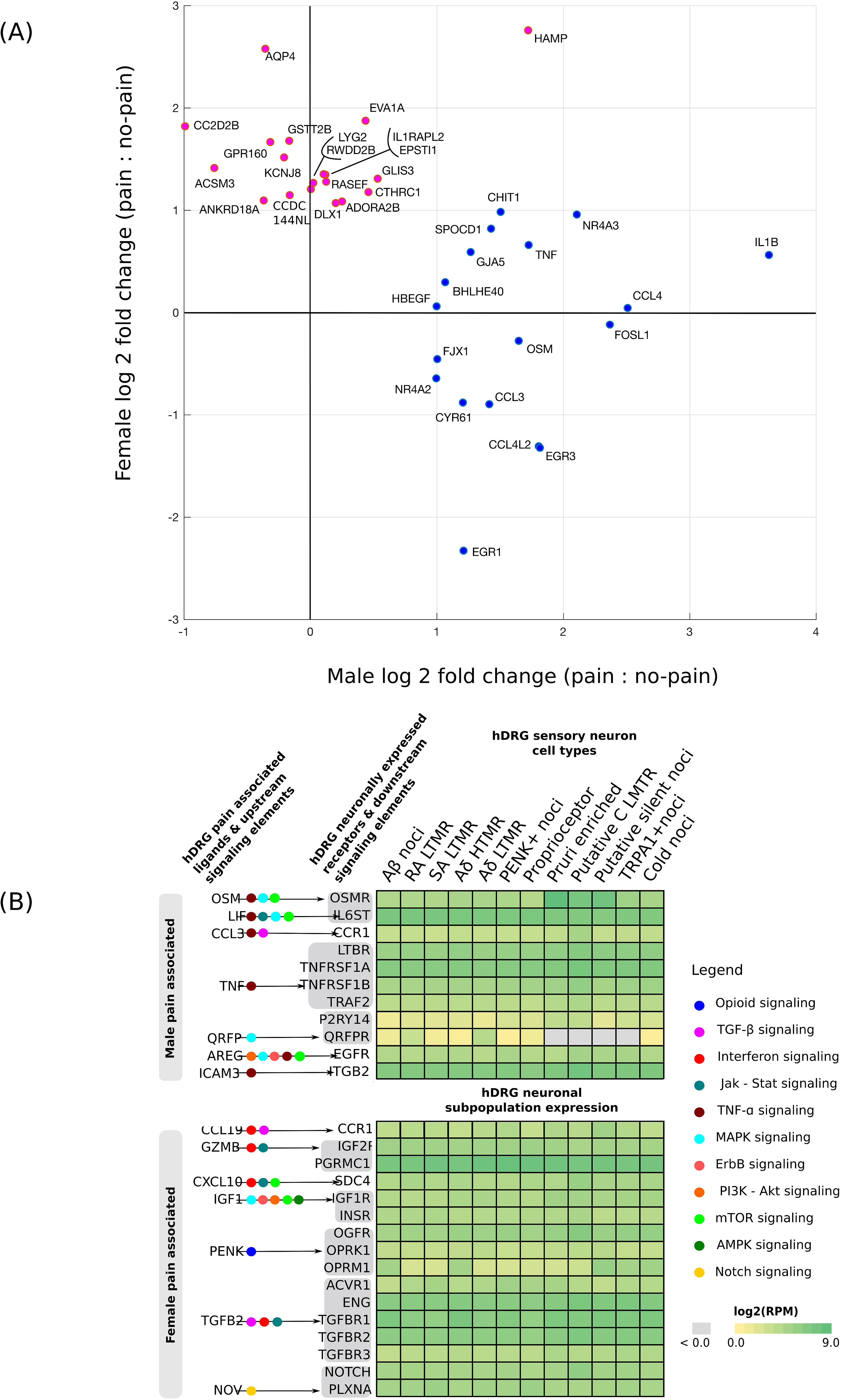
Sex differential aspects of the pain-associated transcriptome. **(A)** The top 25 pain-associated genes (**Supplemental File 3, Sheets A and C**) increased in pain (with a median fold change of 2-fold or higher) for each sex show a remarkably sex differential signal, with only *HAMP* showing 2-fold or greater change in the median in both sexes among the top 25. **(B)** Pain-associated ligands in each sex signal to hDRG-expressed receptors that are enriched in sensory neuronal subpopulations (Tavares-Ferreira et al., 2021) but have little overlap across sexes. Male signaling in enriched in TNF-alpha pathway, while female signaling is enriched in interferon signaling. Though *ICAM3* (in males) and *TGFB2* or *NOV* (in females) are not in the pain-associated gene lists, they are increased in pain for the corresponding sex at the median or upper quartile levels, and are thus shown here.

Intersectional analysis showed overlap in upstream and downstream signaling components for OSM, AP-1, IL-5, IL-1, TNF-alpha, and NFKB pathways in the male pain-associated genes (**Supplemental File 3, Sheet A)**. The TNF-alpha pathway activates the JUN-FOS family of transcription factors that were also pain-associated in our male data. In turn, multiple pro-inflammatory cytokine genes (including IL1B, OSM) as well as TNF, known to be upregulated by the JUN-FOS regulatory cascade, were increased in male pain samples. JUN-FOS and TP53 regulatory signaling was analyzed in more detail in the co-expression module analysis. Other factors known to upregulate JUN/FOS signaling such as LIF (Kanehisa et al., 2007) were also increased in mRNA abundance for male pain samples suggesting redundant upstream signaling through this important transcriptional regulatory pathway. For genes decreasing in male pain samples, we identify several immune cell surface markers (*CD2*, *CD96*), suggesting that immune cell types that decrease in proportion are represented in this list (**Supplemental File 3, Sheet B)**.

For female pain-associated genes, we found overlap with PIK3C, Interferon signaling, JAK-STAT, TLR and IL-5 pathways (**Supplemental File 3, Sheet C**). While *IL5* itself is de-enriched in female pain samples, IL-5 is typically upstream of interferon pathways, and there is likely overlap in IL-5 and interferon signaling pathways. JAK-STAT signaling is also a primary driver of interferon signaling. We found enrichment of both Type I (alpha) and Type II (gamma) interferon signaling pathway genes (both interferon stimulating like *TLR3* and interferon response like *IFIT1*), which are profiled in more detail in the co-expression module analysis. Female pain-associated genes that decrease in the pain state also have some immune cell markers (*CD300LB*, *CCL4L2*), again suggesting that some immune cells increase in frequency at the expense of others (**Supplemental File 3, Sheet D)**.

The overlap of some signaling pathways in males and females (TGF-beta, and IL-5 pathways), but relatively few overlaps in the gene set, suggests that the pain-associated signaling can use common pathways in males and females. However, our findings show that transcriptome enrichment is seen in different parts of the pathway in each sex, similar to our findings in rodent models. This could potentially occur due to sex-specific factors like sex hormone regulation and X-linked gene expression. Additionally, such sex-dimorphic gene expression is likely to drive sex-differential signaling (shown in Figure 3B), as detailed in the following sections.

### Differential expression analysis of paired samples in patients experiencing unilateral pain

We expanded paired analysis for DRGs taken from both sides in male patients with unilateral pain based on our original study (North et al., 2019). We re-analyzed our data with the paired samples from the additional patient and were able to identify 216 genes that increase and 154 genes that decrease in the pain-associated DRGs. These are in close agreement with the distributional analysis-based gene sets identified for the male sub-cohort in the previous section. For instance, we again found a strong enrichment for JUN-FOS signaling in the gene set increased with pain with multiple members of the transcriptional regulatory cascade being increased in abundance (*FOS, FOSB, FOSL1, FOSL2, JUNB, ATF3, EGR1, EGR3*). While JUN-FOS signaling is involved in cellular plasticity in many cell types, it has been implicated in pain since the 1990s (Naranjo et al., 1991) and has recently been implicated as a key regulator of long-term neuropathic pain in mice (Marvaldi et al., 2020). A complete list of the genes increased or decreased in pain samples of matched pairs are presented in **Supplemental File 3, Sheets E and F**. We did not have patient-matched pairs of pain and no-pain DRGs from female patients to complete a similar analysis.

### Mining large co-expressional modules in male and female data for functional annotation

We identified many co-expression modules where gene expression changes were correlated in a sex-stratified sub-cohort. Our work identifies several large co-expression modules – two of which we profiled here: one in males, and one in females. The co-expression module members are listed in **Supplemental File 3, Sheets G - J**, and visualized in Figures 4 **and** 5. We chose to highlight the *OSM* co-expression module in males and *IFIT1* co-expression module in females since they have a large number of co-expressed genes (> 500), well-known connection to pain pathways, and clear evidence of transcriptional regulation (JUN-FOS based regulation in *OSM* module, JAK-STAT signaling in *IFIT1* module).

**Figure 4.**
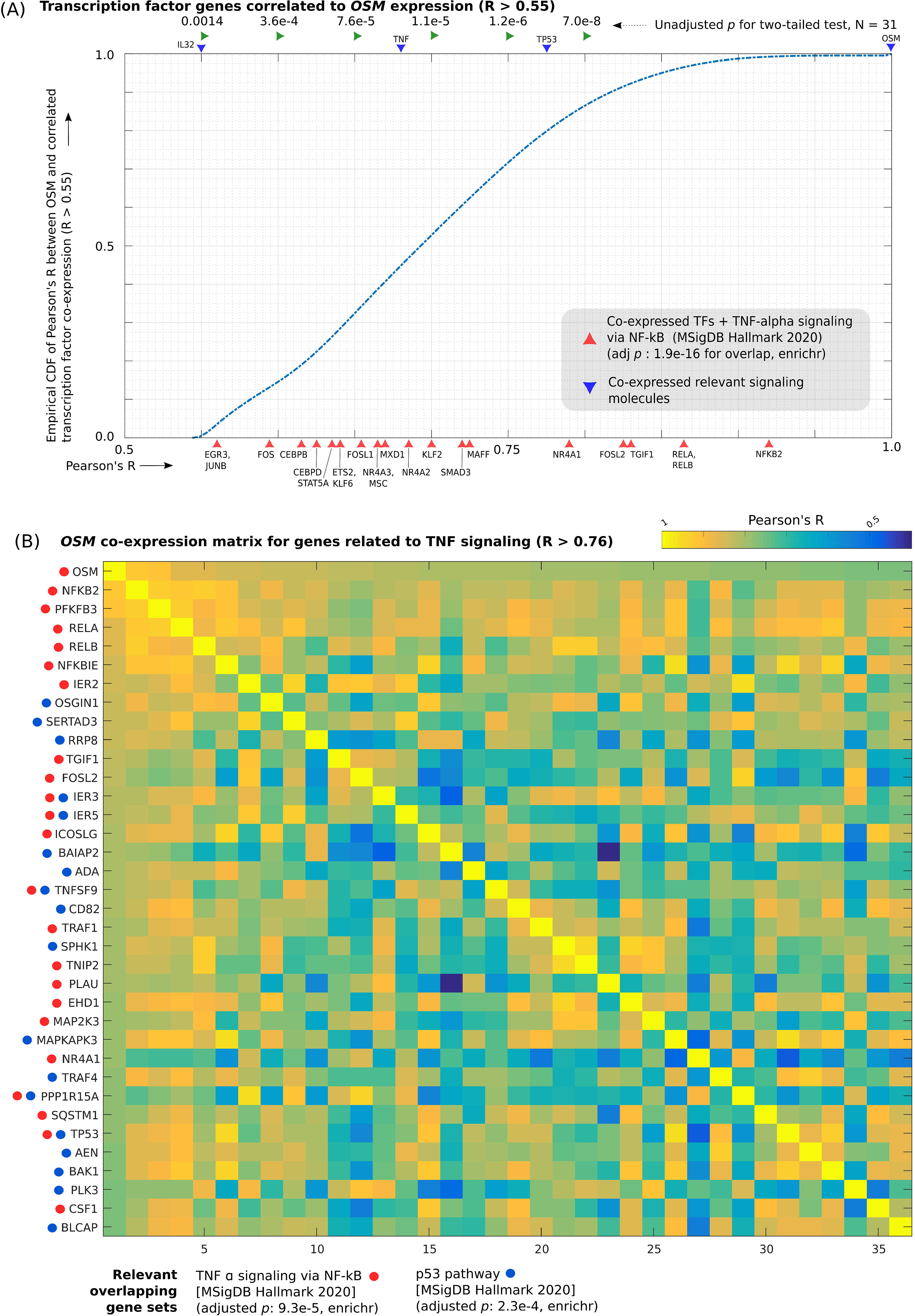
*OSM* co-expression module. **(A)** Empirical CDF of Pearson’s R (with *OSM* expression) for transcription factor genes with Pearson’s R > 0.55, overlaid with Pearson’s R for genes of key signaling molecules in the TNF-alpha pathway (*TNF, TP53* and *OSM*) and *IL32* (another pro-inflammatory cytokine) and members of enriched gene sets that overlap with the co-expressed TF genes. **(B)** Heatmap of correlation matrix (Pearson’s R) between members of *OSM* co-expression module in enriched gene sets (TNF signaling via NFKB, p53 signaling) that overlap with it.

**Figure 5.**
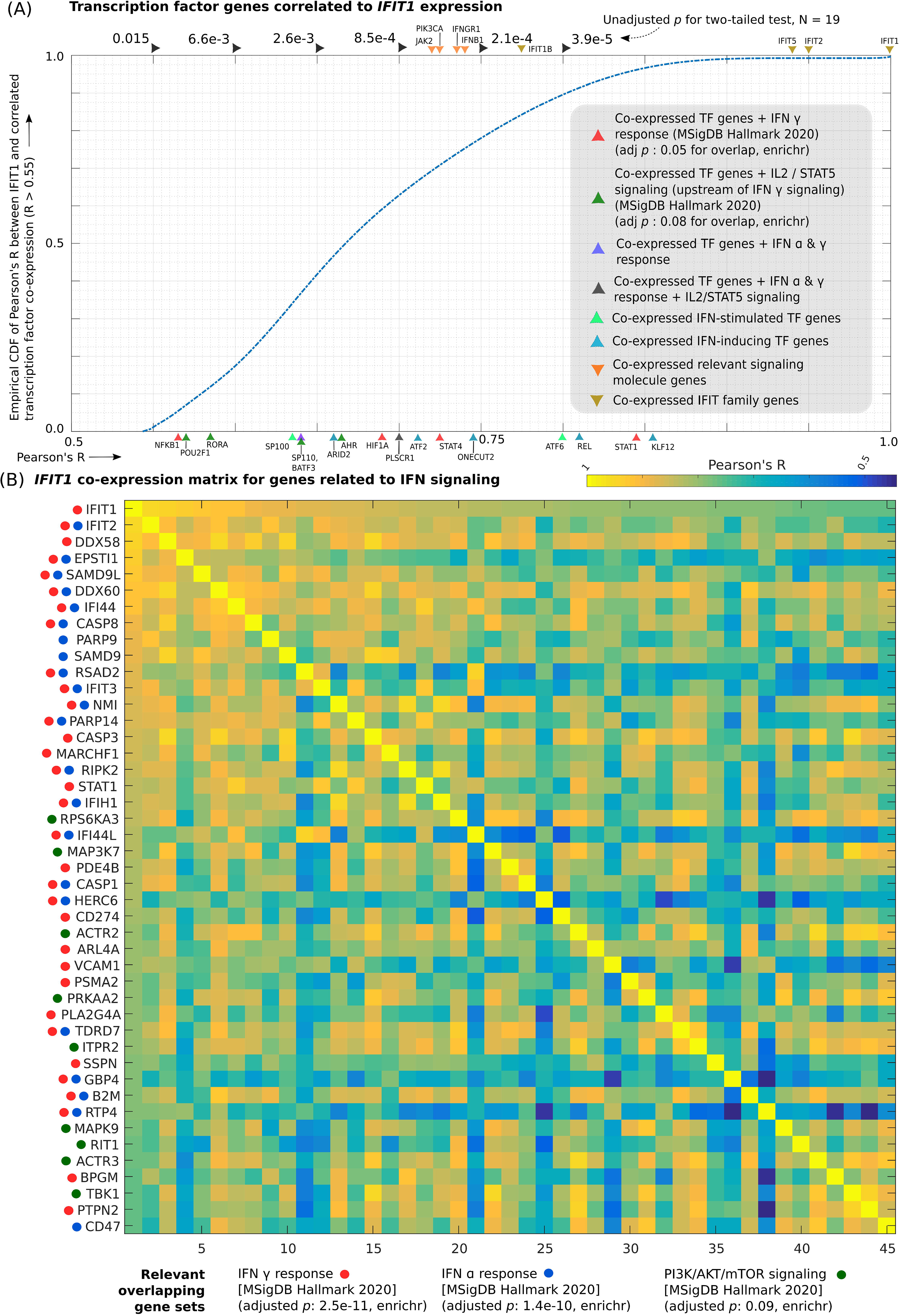
*IFIT1* co-expression module. **(A)** Empirical CDF of Pearson’s R (with *IFIT1* expression) for transcription factor genes with Pearson’s R > 0.55, overlaid with Pearson’s R for genes of key signaling molecules in the interferon signaling pathways and members of enriched gene sets that overlap with the co-expressed TF genes. **(B)** Heatmap of correlation matrix (Pearson’s R) between members of *IFIT1* co-expression module in enriched gene sets (interferon alpha and gamma response, PI3K/AKT/mTOR signaling) that overlap with it.

We identified 209 TF genes that were correlated in expression to *OSM* (**Supplemental File 3, Sheet G)** including several that are well known to be implicated in the TNF and OSM pathways (several members of the JUN-FOS family, *RELA, RELB, NFKB2, CEBPB, TP53* shown in Figure 4A). We found that the *OSM* co-expression module included 697 genes, including pro-inflammatory cytokines such as *OSM* and *CXCL16,* MAPK signaling genes (*MAP2K3, MAPKAPK3*), TP53 pathway (*TP53, PLK3, TRAF4*) and TNF-signaling (PLAU, TNFSF9, CSF1) pathway molecules and was the largest pain-associated co-expression module in males **(Supplemental File 3, Sheet H and** Figure 4B). Multiple members of the JUN-FOS signaling cascade (*FOS, FOSB, JUNB, EGR3*) that are regulated by the JUN-FOS (AP-1) transcriptional regulation and TNF signaling were present in this module. *TNF* itself and pro-inflammatory cytokines like *IL32* were co-expressed with OSM, though with a correlation coefficient between 0.55 and 0.76, likely because they are upstream of OSM or co-regulated (Figure 4A). This finding suggests a key role for active transcriptional programs in pro-inflammatory cytokine signaling within the DRG in neuropathic pain. In rodent models, importin α3 has been shown to be crucial for nuclear import of FOS, known to be important in maintenance of chronic pain (Marvaldi et al., 2020; Yousuf and Price, 2020) suggesting that a similar mechanism may be at work in humans.

We found 130 TF genes to be co-expressed with *IFIT1*, including multiple TF genes that are interferon-inducing (*KLF12, REL, ONECUT2*) or interferon-stimulated (*STAT1, STAT4, RORA, NFKB1*), suggesting a role for interferon signaling in this pathway **(Supplemental File 3, Sheet I and** Figure 5A). Also, we found 743 genes to be co-expressed with IFIT1 across the whole transcriptome including *IFIT1* and *IFIT2,* interferon-induced proteins that are involved in autoimmune disorders like Lupus (Ye et al., 2003) **(Supplemental File 2, Sheet J and** Figure 5B). Other genes of the same family *IFIT1B*, *IFIT3*, and *IFIT5* were part of the same module. STAT1, a transcription factor that is part of the JAK-STAT signaling pathway, a key component of interferon signaling. (Schindler et al., 2007) is present. Multiple other interferon-induced genes like *IFI44*, *TLR1*, *CASP3*, and *CASP8* suggest increased interferon signaling in females with neuropathic pain, possibly in immune cells. Genes like *KLF12* and *REL* suggest a role of T-cells in the identified interferon signaling pathway.

### Protein interaction networks

Many of the pain-associated genes we identified were immune signaling or immune response genes, consistent with a neuro-immune signaling complex in neuropathic pain DRGs that may be driving changes in nociceptor excitability. We found multiple ligand genes with increased abundance in pain that can signal to neuronally-expressed receptors (Wangzhou et al., 2021) (Figure 3B) enriched or expressed in hDRG neuronal subpopulations (that we described previously using spatial transcriptomics (Tavares-Ferreira et al., 2021)). An example is *QRFP*, which is increased in male pain, co-expressed with *OSM* and signals to the hDRG specific *QRFPR* that encodes a receptor signaling through the mTOR pathway (Li et al., 2021), a pathway that is critical for regulation of nociceptor excitability (Jimenez-Diaz et al., 2008; Geranton et al., 2009; Melemedjian et al., 2011; Megat et al., 2019). While multiple pathways are possibly involved in signaling from immune cells with pro-inflammatory phenotypes to sensory neurons, we found that TNF-alpha signaling components were present in multiple putative ligand – receptor interactions, where the ligand was male pain-associated (*OSM, LIF, CCL3, TNF, AREG, ICAM3*), and the receptor was known to be expressed in hDRG sensory neurons. In female samples, multiple interferon signaling related ligand - receptor interactions were identified where the ligand was female pain-associated, and the receptor was expressed in human sensory neurons (*CCL19, GZMB, CXCL10, IGF1*).

StringDB-based protein interaction networks for the male and female pain-associated genes are presented in **Supplementary Figures 1 and 2**. In the male signaling network (**Supplementary Figure 1**), a densely connected network of 9 genes (including *TNF*), which are expressed in the hDRG interact with pain-associated gene products. More importantly, members of this network, as well as *IL1B* and *CCL4* are part of multiple, interwoven signaling networks (TNF, TLR, growth factor, and cytokine-based signaling) associated with neuropathic pain in males.

In the female signaling network (**Supplementary Figure 2**), we identified multiple type I and II interferon-stimulated genes *(IFIT1/2, GBP2/3/7, CD2, HLA-DQB1, CXCL10*). This again supports a prominent role of interferon signaling in neuropathic pain in females. The role of genes like *IFIT1*, and *IFIT2* in diseases like Lupus, and Sickle Cell Disease (Ye et al., 2003; Hermand-Tournamille et al., 2018) and the role of female sex hormones in interferon signaling in autoimmune diseases (Singh and Hahn, 2020) suggest the possibility of autoimmune signaling in a subset of the female pain samples.

Additionally, densely connected subgraphs for centromeric proteins (**Supplementary Figure 2**), and proteins involved in cell cycle and proliferation suggest that a significant part of the bulk RNA-seq transcriptomes are driven by proliferating cell types in both sexes. These cells are likely pro-inflammatory because in both the male and female signaling networks inflammatory cytokine signaling was identified.

### RNAscope analysis

Based on our findings from the RNA-seq analysis, we performed RNAscope *in situ* hybridization on a subset of pain and non-pain samples to assess the cellular distribution of differentially expressed genes. We used a qualitative approach due to the limited availability of banked fresh-frozen tissue samples. For RNAscope, besides our genes of interest, we used the additional channels for marker genes to label specific cell types. We used *TRPV1* to label all nociceptors (Shiers et al., 2020; Shiers et al., 2021; Tavares-Ferreira et al., 2021), *AIF1*, which is expressed by the monocyte / macrophage lineages in the nervous system (Schwab et al., 2001), as well as some T-cells (Galdo and Jimenez, 2007), and DAPI to label all nuclei.

#### RNAscope for male samples

For the male samples, we examined cellular expression profiles of several genes that were well correlated with *OSM* expression (all with Pearson’s R >= 0.54, uncorrected *p* <= 0.0017): *TNF, IL32, IL1B* and *OSM* itself. OSM signals through the GP-130 complex, using OSMR as a co-receptor (Taga, 1996). OSM is known to promote nociceptor sensitization in rodents (Langeslag et al., 2011; Garza Carbajal et al., 2021) suggesting that it may be a key signaling molecule for neuropathic pain in humans, in particular males. Our RNAscope imaging showed *TNF* expression in both nociceptors (TRPV1+ cells) and some *TRPV1*-sensory neurons, *AIF*+ immune cells and some *AIF*- non-neuronal cells, suggesting that immune cell driven TNF signaling could promote TNF expression in nociceptors, or vice versa, in a feedback loop (Figure 6A and B). Additionally, we found *OSM*, *IL32* and *IL1B* primarily expressed in *AIF1*+ immune cells, likely macrophages, but we additionally note some expression in *AIF*- non-neuronal cells (Figure 6C and D). Qualitatively, transcript abundance (puncta) increased in all of these genes in the pain samples, compared to the non-pain control, supporting the transcriptomic findings. One of the pain samples (84T4L) had considerably more signal for many of these genes.

**Figure 6.**
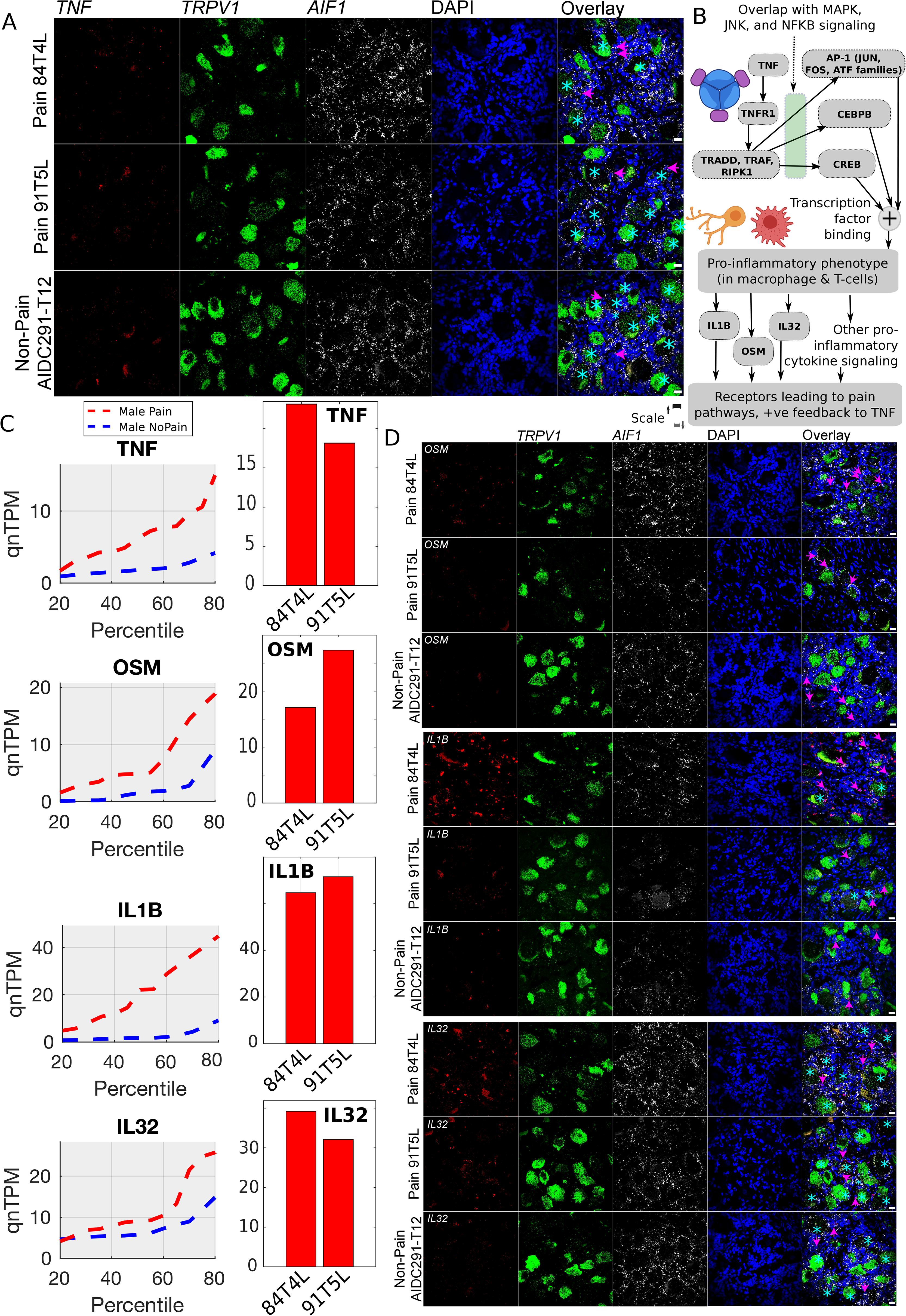
RNAscope for male pain-associated genes *TNF, OSM, IL1B,* and *IL32*. **(A)** RNAscope *TNF* expression (red) overlaid with *TRPV1* expression (green), *AIF1* (white) and DAPI (blue) in pseudocolor. **(B)** Schema showing how TNF-alpha signaling cascades overlap with other signaling pathways, based on existing literature (icons from BioRender) **(C**) *TNF, OSM, IL1B* and *IL32* quantile plots show the shift in abundance between male pain and non-pain sub-cohorts, while the bar plots show the RNA-seq abundance for the patient DRGs that were queried by RNAscope assay **(D)** RNAscope *OSM, IL1B, and IL32* expression (red) overlaid with *TRPV1* expression (green), *AIF1* (white) and DAPI (blue) in pseudocolor. Scalebar = 20 μm, large globular structures are considered to be lipofuscin. Arrows point to target-positive (red) signal in non-neuronal cells, and stars denote target-positive signal (red) in neurons. (**A** and **D**)

#### RNAscope for female samples

Among the female pain-associated genes we found several (*IFIT1, IFIT2, HLA-DQB1, TLR1, CXCL10, NMI*) that could be type I or II interferon-stimulated and had a high dynamic range of 10-fold or higher variance across female pain samples (Figure 7A-E). This supports the conclusion that a subset of female pain samples had activated interferon signaling pathways (Figure 7D). Consistent with this, we found qualitatively more *IFIT1* expression in the female pain samples. The gene was found in *TRPV1*-positive and *TRPV1*-negative neurons but showed the highest abundance in non-neuronal cells that were both *AIF1*-positive and *AIF1*-negative (Figure 7A and B). *HLA-DQB1* was highly expressed ubiquitously, though the very high expression of *HLA-DQB1* in 86T6R RNA-seq was not qualitatively observed in the RNAscope assay (Figure 7B and C).

**Figure 7.**
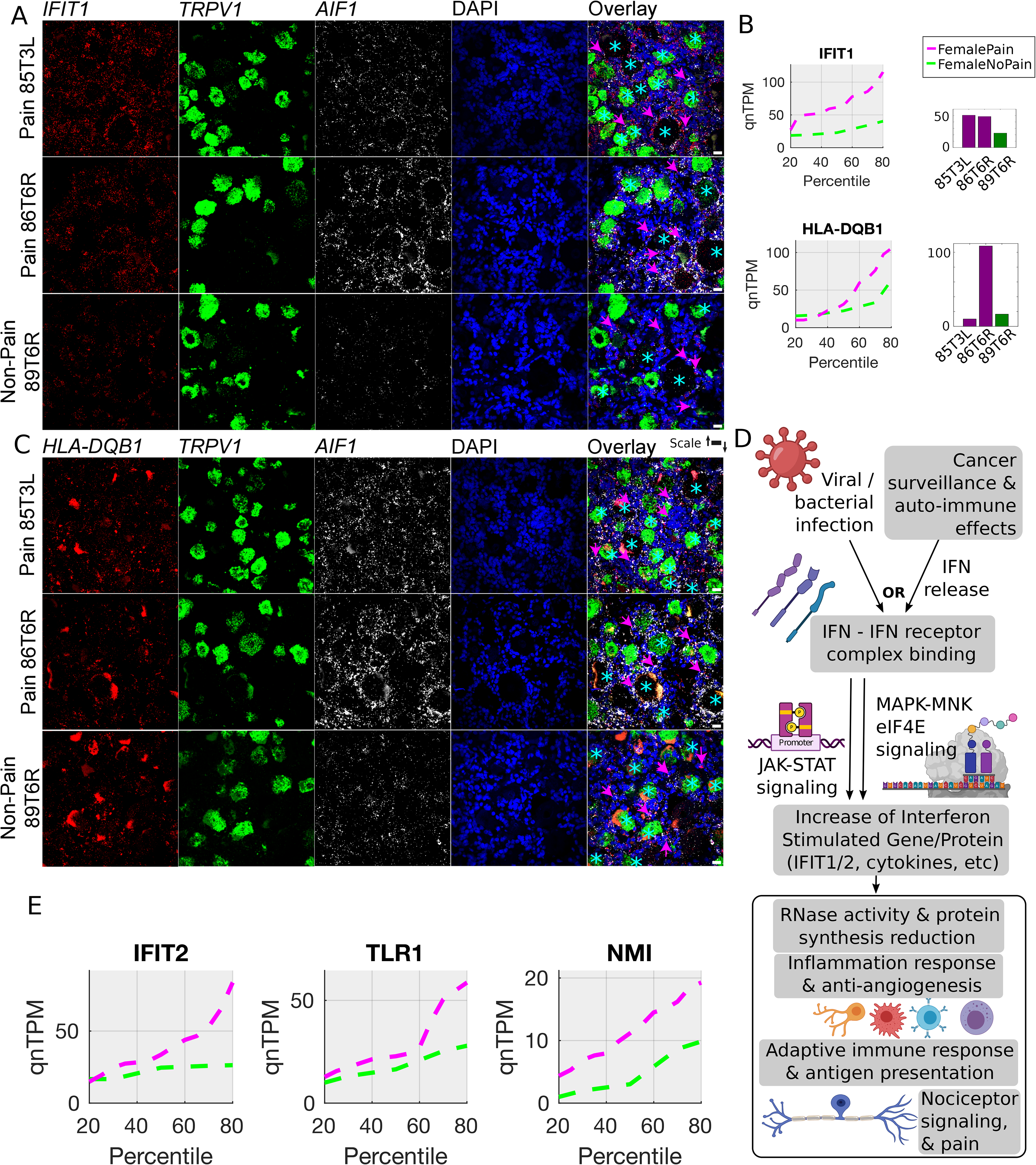
RNAscope for female pain-associated genes *IFIT1* and *HLA-DQB1*. **(A)** RNAscope *IFIT1* expression (red) overlaid with *TRPV1* expression (green), *AIF1* (white) and DAPI (blue) in pseudocolor. **(B**) *IFIT1* and *HLA-DQB1* quantile plots show the shift in abundance between female pain and non-pain sub-cohorts, while the bar plots show the RNA-seq abundance for the patient DRGs that were queried by RNAscope assay **(C)** RNAscope *HLA-DQB1* expression (red) overlaid with *TRPV1* expression (green), *AIF1* (white) and DAPI (blue) in pseudocolor. **(D)** Schema showing how interferons are typically activated and drive a signaling program, based on existing literature (icons from BioRender) **(E)** Quantile plots in the female cohort for other *IFIT1*-correlated interferon signaling pathway genes - *IFIT2, TLR1, NMI -* show increase in gene abundance in pain. Scalebar = 20 μm, large globular structures are considered to be lipofuscin. Arrows point to target-positive (red) signal in non-neuronal cells, and stars denote target-positive signal (red) in neurons. (**A** and **C**)

A consistent observation from the female RNAscope assays was the high abundance of *AIF1*-signal in the female pain samples (85T3L and 86T6R), suggesting either macrophage infiltration into the hDRG or an increase in *AIF1* transcription due to macrophage activation (Figure 7A and C). Our images suggest the latter as we observe no visible difference in cell numbers; however, a formal analysis with more biological replicates would be needed to confirm this observation.

These cell-type specific expression studies provide consistent evidence of increased *AIF1*+ cell expression in pain-associated samples in males and females. However, these macrophages or T-cells likely have different expression patterns in men and women given the upregulation of specific gene sets that we found to be enriched in this cell population using RNAscope. Our findings are consistent with emerging literature using rodents where macrophages and T cells are key players in neuropathic pain but with different signaling molecules from these cells promoting pain depending on sex (Liang et al., 2020; Yu et al., 2020; Rudjito et al., 2021).

## Discussion

We reach several conclusions based on the data presented here. First, we observed major sex differences in changes in gene expression in the human DRG associated with neuropathic pain. This difference is consistent with a growing number of findings from rodent models (reviewed in (Mogil, 2020)), but our findings highlight different sets of genes as potential drivers of neuropathic pain, in particular in females. Second, our work highlights the important role that neuro-immune interactions likely play in causing neuropathic pain (Ji et al., 2016). These cell types may differ between men and women (Sorge et al., 2015; Yu et al., 2020; Rudjito et al., 2021), and the ligand-receptor interactions that allow these cells to communicate with nociceptors in the DRG almost certainly differ between men and women, even if cell types are consistent between sexes. Finally, our work points to therapeutic targets for neuropathic pain based on molecular neuroscience insight in patients. Again, these targets are sex dimorphic with a FOS/JUN-driven cytokine profile a dominant feature in males and type I and II interferon signaling playing a key role in females. Additional work will be needed to evaluate the clinical translatability of these findings.

One potential shortcoming of our work is the use of bulk rather than single cell or spatial transcriptomics. These single cell (Nguyen et al., 2021) and spatial (Tavares-Ferreira et al., 2021) technologies have now been used successfully on human DRG, hence these technologies will be valuable for future studies on human DRG samples from thoracic vertebrectomy patients. We chose to focus on bulk sequencing because we did not have an *a priori* notion of which cell types to focus on and hence were unable to do targeted sequencing of specific cell types, and we were concerned about financial constraints on the large sample size required to thoroughly assess sex differences in this patient population. In that regard, this study builds upon our previously published work where we speculated on sex differences in neuropathic pain-associated expression changes in the DRGs of thoracic vertebrectomy patients (North et al., 2019), but we did not have a sufficient sample size to make such a direct comparison in that study. We have now achieved a sufficient sample size and observed robust changes in gene expression associated with neuropathic pain in both sexes. This justifies the choice of bulk sequencing and shows that it gives a unique insight into overall changes in gene expression that can now be exploited more thoroughly using single cell and spatial techniques. Our work also generates a set of testable hypotheses, e.g. the effect of interferons on female DRG neurons, that can be tested in physiological experiments. Finally, RNAscope examines cellular expression patterns and shows that changes in gene expression can be observed across multiple cell types demonstrating that spatial transcriptomics approaches may be the most suitable for understanding the complex interactions that likely occur between non-neuronal and neuronal cells in the neuropathic DRG. Our future work will employ this technology to better understand neuro-immune interactions and the immune and neuronal subpopulations involved, as we have already demonstrated its utility in identifying sensory neuron subtypes in DRGs from organ donors (Tavares-Ferreira et al., 2021).

Spontaneous or ectopic activity in DRG neurons is a likely cause of neuropathic pain (Haroutounian et al., 2014; Vaso et al., 2014; North et al., 2019; Middleton et al., 2021) and may be a driver of pain in other diseases, like fibromyalgia (Serra et al., 2014). This ectopic activity generates an aberrant signal that is conveyed through the spinal cord to the brain to cause pain sensations in people who suffer from neuropathic pain. Ectopic activity can originate from axons, in particular early after injury, but it can also emerge from the DRG soma (Amir et al., 2005). This likely happens due to changes in expression of ion channels in the DRG neuronal soma (Devor, 2006), or due to signaling events that cause the neuron to generate ectopic action potentials and instability in the resting membrane potential (Devor and Yarom, 2002; Odem et al., 2018; Garza Carbajal et al., 2020; Laumet et al., 2020; Lopez et al., 2021). Our previous work clearly showed that human DRG neuron somata from neuropathic pain patients could display ectopic activity, even after surgical removal and culturing (North et al., 2019). Our current work gives important clues to what might drive these changes in patients. Moreover, our findings suggest that although the resultant DRG neuronal phenotype in males and females is the same, ectopic activity, it is likely driven by unique intercellular mechanisms. In males those mechanisms appear to involve cytokines like TNFα, IL1β and OSM that likely originate from macrophages, but may also be released from neurons themselves. These findings in males are supported by findings from animal models where macrophages have been implicated in development of chronic pain (Yu et al., 2020; Rudjito et al., 2021). In females the picture is much different. We discovered a network of genes associated with type I and II interferons that are positively correlated with neuropathic pain in women undergoing thoracic vertebrectomy. Interferons have been linked to pain sensitization (Barragan-Iglesias et al., 2020), and also to analgesia (Donnelly et al., 2021), by previous studies but their sex-specific role in neuropathic pain has not been studied. Examining the effect of type I and II interferons on female human DRG neurons should be a priority for future work.

While our work gives unique insight into mechanisms of neuropathic pain, it is not feasible to sample the DRG of the vast majority of pain patients. How can this work be used to facilitate precision medicine that can address underlying pain mechanisms in individual patients? Our findings demonstrate that immune cells in the DRG take on a proliferative role in neuropathic pain and these cells may be infiltrating the DRG from the peripheral circulation. It is also possible that the cytokines generated within the DRG may have an influence on peripheral immune cells. Both scenarios raise the possibility that specific cell types in the blood could be sampled to gain insight into molecular changes in the DRG without actually accessing the DRG. Animal studies have demonstrated that brain transcriptome changes in neuropathic pain models can be represented in immune cells transcriptomes in the periphery (Massart et al., 2016). Likewise, clinical studies have shown blood transcriptomic differences that predict chronic low back pain susceptibility (Dorsey et al., 2019). A key question is which cells to sample to gain the greatest insight into underlying mechanisms. From our work we propose that the candidate cell types may differ by sex. In males, monocytes would be a clear candidate cell type given the changes observed in male neuropathic pain DRGs. In females this is less clear, but T-cells and monocytes would be good candidates. One possibility is that monocytes are key players in both sexes, but these cells simply have fundamentally different repertoires of ligands that they express that then communicate with unique receptors expressed by DRG neurons. They may also have unique honing mechanisms that drive cells from the peripheral circulation into the DRG.

In closing, our findings make a strong case for important sex differences in neuropathic pain mechanisms in the human DRG. Male mechanisms are closely tied to inflammatory cytokines that have been studied widely in preclinical models for decades. Female mechanisms appear to involve interferon stimulated genes and represent signaling mechanisms that are not widely studied in preclinical models. This highlights the importance of considering sex as a biological variable, not only for basic science insight, but also in developing therapeutic strategies to treat neuropathic pain.

## Author contributions

PRR, SS, PMD and TJP conceived and designed the overall study. RYN and CT recruited patients and did the surgeries. MLU and YL recovered tissues after surgery and did tissue processing. DTF designed and performed the experiments for the Visium neuronal transcriptome profiles. PRR designed and performed all bioinformatic analyses, with the exception of the Visium neuronal transcriptome profiles. SS designed and performed all RNAscope analyses. IS, GD and MB provided input on immune cell lineages and markers. PRR, SS and TJP wrote the manuscript, with input from GD. PMD and TJP supervised the study. All authors edited the manuscript and agreed to the final version.

## Acknowledgements

The authors would like to thank all the tissue donor patients and their family, and the surgical and medical staff without whose assistance this work would not be possible. The authors would also like to thank all members of the genome sequencing core at UT Dallas, and Andi Wangzhou for help with the sequencing and mapping protocols.

## Funding

NS111929 (to PMD and TJP), NS102161 (to TJP)

## Competing interests

The authors declare no competing interests for this work. TJP, GD and PRR are co-founders of Doloromics.

## Notes

https://utdallas.box.com/s/7uwuqr1pgp268je7vnae9uao624owp14

## References

Agalave NM, Rudjito R, Farinotti AB, Khoonsari PE, Sandor K, Nomura Y, Szabo-Pardi TA, Urbina CM, Palada V, Price TJ, Erlandsson Harris H, Burton MD, Kultima K, Svensson CI (2021) Sex-dependent role of microglia in disulfide high mobility group box 1 protein-mediated mechanical hypersensitivity. Pain 162:446–458.

Amir R, Kocsis JD, Devor M (2005) Multiple interacting sites of ectopic spike electrogenesis in primary sensory neurons. J Neurosci 25:2576–2585.

Anders S, Huber W (2010) Differential expression analysis for sequence count data. Nature Precedings:1–1.

Barragan-Iglesias P, Franco-Enzastiga U, Jeevakumar V, Shiers S, Wangzhou A, Granados-Soto V, Campbell ZT, Dussor G, Price TJ (2020) Type I Interferons Act Directly on Nociceptors to Produce Pain Sensitization: Implications for Viral Infection-Induced Pain. J Neurosci 40:3517–3532.

Dahlhamer J, Lucas J, Zelaya C, Nahin R, Mackey S, DeBar L, Kerns R, Von Korff M, Porter L, Helmick C (2018) Prevalence of chronic pain and high-impact chronic pain among adults—United States, 2016. Morbidity and Mortality Weekly Report 67:1001.

Davidson S, Golden JP, Copits BA, Ray PR, Vogt SK, Valtcheva MV, Schmidt RE, Ghetti A, Price TJ, Gereau RWt (2016) Group II mGluRs suppress hyperexcitability in mouse and human nociceptors. Pain 157:2081–2088.

Devor A, Yarom Y (2002) Generation and propagation of subthreshold waves in a network of inferior olivary neurons. J Neurophysiol 87:3059–3069.

Devor M (2006) Sodium channels and mechanisms of neuropathic pain. J Pain 7:S3–S12.

Dobin A, Davis CA, Schlesinger F, Drenkow J, Zaleski C, Jha S, Batut P, Chaisson M, Gingeras TR (2013) STAR: ultrafast universal RNA-seq aligner. Bioinformatics 29:15–21.

Donnelly CR, Jiang C, Andriessen AS, Wang K, Wang Z, Ding H, Zhao J, Luo X, Lee MS, Lei YL, Maixner W, Ko MC, Ji RR (2021) STING controls nociception via type I interferon signalling in sensory neurons. Nature 591:275–280.

Dorsey SG, Renn CL, Griffioen M, Lassiter CB, Zhu S, Huot-Creasy H, McCracken C, Mahurkar A, Shetty AC, Jackson-Cook CK, Kim H, Henderson WA, Saligan L, Gill J, Colloca L, Lyon DE, Starkweather AR (2019) Whole blood transcriptomic profiles can differentiate vulnerability to chronic low back pain. PLoS One 14:e0216539.

Finnerup NB, Attal N, Haroutounian S, McNicol E, Baron R, Dworkin RH, Gilron I, Haanpaa M, Hansson P, Jensen TS, Kamerman PR, Lund K, Moore A, Raja SN, Rice AS, Rowbotham M, Sena E, Siddall P, Smith BH, Wallace M (2015) Pharmacotherapy for neuropathic pain in adults: a systematic review and meta-analysis. Lancet Neurol 14:162–173.

Frankish A, Diekhans M, Ferreira A-M, Johnson R, Jungreis I, Loveland J, Mudge JM, Sisu C, Wright J, Armstrong J (2019) GENCODE reference annotation for the human and mouse genomes. Nucleic acids research 47:D766–D773.

Galdo FD, Jimenez SA (2007) T cells expressing allograft inflammatory factor 1 display increased chemotaxis and induce a profibrotic phenotype in normal fibroblasts in vitro. Arthritis & Rheumatism: Official Journal of the American College of Rheumatology 56:3478–3488.

Garza Carbajal A, Bavencoffe A, Walters ET, Dessauer CW (2020) Depolarization-Dependent C-Raf Signaling Promotes Hyperexcitability and Reduces Opioid Sensitivity of Isolated Nociceptors after Spinal Cord Injury. J Neurosci 40:6522–6535.

Garza Carbajal A, Ebersberger A, Thiel A, Ferrari L, Acuna J, Brosig S, Isensee J, Moeller K, Siobal M, Rose-John S, Levine J, Schaible HG, Hucho T (2021) Oncostatin M induces hyperalgesic priming and amplifies signaling of cAMP to ERK by RapGEF2 and PKA. J Neurochem 157:1821–1837.

Geranton SM, Jimenez-Diaz L, Torsney C, Tochiki KK, Stuart SA, Leith JL, Lumb BM, Hunt SP (2009) A rapamycin-sensitive signaling pathway is essential for the full expression of persistent pain states. J Neurosci 29:15017–15027.

Haanpää M, Attal N, Backonja M, Baron R, Bennett M, Bouhassira D, Cruccu G, Hansson P, Haythornthwaite JA, Iannetti GD (2011) NeuPSIG guidelines on neuropathic pain assessment. PAIN® 152:14–27.

Haroutounian S, Nikolajsen L, Bendtsen TF, Finnerup NB, Kristensen AD, Hasselstrom JB, Jensen TS (2014) Primary afferent input critical for maintaining spontaneous pain in peripheral neuropathy. Pain 155:1272–1279.

Hermand-Tournamille P, Azouzi S, Salnot V, Gautier EF, Mayeux P, Colin Y, Bondet V, Duffy D, Tharaux PL, Kim CLV (2018) Proteomic landscape of neutrophils in sickle cell anemia: an unexpected autoimmune profile. Blood 132:2357.

Ho JW, Stefani M, Dos Remedios CG, Charleston MA (2008) Differential variability analysis of gene expression and its application to human diseases. Bioinformatics 24:i390–i398.

Inyang KE, Szabo-Pardi T, Wentworth E, McDougal TA, Dussor G, Burton MD, Price TJ (2019) The antidiabetic drug metformin prevents and reverses neuropathic pain and spinal cord microglial activation in male but not female mice. Pharmacol Res 139:1–16.

Jeanquartier F, Jean-Quartier C, Holzinger A (2015) Integrated web visualizations for protein-protein interaction databases. BMC Bioinformatics 16:195.

Jensen TS, Baron R, Haanpää M, Kalso E, Loeser JD, Rice AS, Treede R-D (2011) A new definition of neuropathic pain. Pain 152:2204–2205.

Ji RR, Chamessian A, Zhang YQ (2016) Pain regulation by non-neuronal cells and inflammation. Science 354:572–577.

Jimenez-Diaz L, Geranton SM, Passmore GM, Leith JL, Fisher AS, Berliocchi L, Sivasubramaniam AK, Sheasby A, Lumb BM, Hunt SP (2008) Local translation in primary afferent fibers regulates nociception. PLoS One 3:e1961.

Kanehisa M, Araki M, Goto S, Hattori M, Hirakawa M, Itoh M, Katayama T, Kawashima S, Okuda S, Tokimatsu T (2007) KEGG for linking genomes to life and the environment. Nucleic acids research 36:D480–D484.

Kuleshov MV, Jones MR, Rouillard AD, Fernandez NF, Duan Q, Wang Z, Koplev S, Jenkins SL, Jagodnik KM, Lachmann A, McDermott MG, Monteiro CD, Gundersen GW, Ma’ayan A (2016) Enrichr: a comprehensive gene set enrichment analysis web server 2016 update. Nucleic Acids Res 44:W90–97.

LaCroix-Fralish ML, Austin J-S, Zheng FY, Levitin DJ, Mogil JS (2011) Patterns of pain: meta-analysis of microarray studies of pain. PAIN® 152:1888–1898.

Lambert SA, Jolma A, Campitelli LF, Das PK, Yin Y, Albu M, Chen X, Taipale J, Hughes TR, Weirauch MT (2018) The Human Transcription Factors. Cell 172:650–665.

Langeslag M, Constantin CE, Andratsch M, Quarta S, Mair N, Kress M (2011) Oncostatin M induces heat hypersensitivity by gp130-dependent sensitization of TRPV1 in sensory neurons. Mol Pain 7:102.

Laumet G, Bavencoffe A, Edralin JD, Huo XJ, Walters ET, Dantzer R, Heijnen CJ, Kavelaars A (2020) Interleukin-10 resolves pain hypersensitivity induced by cisplatin by reversing sensory neuron hyperexcitability. Pain 161:2344–2352.

Li H, Lou R, Xu X, Xu C, Yu Y, Xu Y, Hu L, Xiang Y, Lin X, Tang S (2021) The variations in human orphan G protein-coupled receptor QRFPR affect PI3K-AKT-mTOR signaling. Journal of Clinical Laboratory Analysis:e23822.

Liang Z, Hore Z, Harley P, Stanley FU, Michrowska A, Dahiya M, La Russa F, Jager SE, Villa-Hernandez S, Denk F (2020) A transcriptional toolbox for exploring peripheral neuro-immune interactions. Pain.

Liberzon A, Birger C, Thorvaldsdottir H, Ghandi M, Mesirov JP, Tamayo P (2015) The Molecular Signatures Database (MSigDB) hallmark gene set collection. Cell Syst 1:417–425.

Lopez ER, Carbajal AG, Tian JB, Bavencoffe A, Zhu MX, Dessauer CW, Walters ET (2021) Serotonin enhances depolarizing spontaneous fluctuations, excitability, and ongoing activity in isolated rat DRG neurons via 5-HT4 receptors and cAMP-dependent mechanisms. Neuropharmacology 184:108408.

Marvaldi L, Panayotis N, Alber S, Dagan SY, Okladnikov N, Koppel I, Di Pizio A, Song DA, Tzur Y, Terenzio M, Rishal I, Gordon D, Rother F, Hartmann E, Bader M, Fainzilber M (2020) Importin alpha3 regulates chronic pain pathways in peripheral sensory neurons. Science 369:842–846.

Massart R, Dymov S, Millecamps M, Suderman M, Gregoire S, Koenigs K, Alvarado S, Tajerian M, Stone LS, Szyf M (2016) Overlapping signatures of chronic pain in the DNA methylation landscape of prefrontal cortex and peripheral T cells. Sci Rep 6:19615.

Megat S, Ray PR, Moy JK, Lou TF, Barragan-Iglesias P, Li Y, Pradhan G, Wanghzou A, Ahmad A, Burton MD, North RY, Dougherty PM, Khoutorsky A, Sonenberg N, Webster KR, Dussor G, Campbell ZT, Price TJ (2019) Nociceptor Translational Profiling Reveals the Ragulator-Rag GTPase Complex as a Critical Generator of Neuropathic Pain. J Neurosci 39:393–411.

Melemedjian OK, Asiedu MN, Tillu DV, Sanoja R, Yan J, Lark A, Khoutorsky A, Johnson J, Peebles KA, Lepow T, Sonenberg N, Dussor G, Price TJ (2011) Targeting adenosine monophosphate-activated protein kinase (AMPK) in preclinical models reveals a potential mechanism for the treatment of neuropathic pain. Mol Pain 7:70.

Menashe I, Aloni R, Lancet D (2006) A probabilistic classifier for olfactory receptor pseudogenes. BMC Bioinformatics 7:393.

Middleton SJ, Barry AM, Comini M, Li Y, Ray PR, Shiers S, Themistocleous AC, Uhelski ML, Yang X, Dougherty PM, Price TJ, Bennett DL (2021) Studying human nociceptors: from fundamentals to clinic. Brain 144:1312–1335.

Mogil JS (2020) Qualitative sex differences in pain processing: emerging evidence of a biased literature. Nat Rev Neurosci 21:353–365.

Naranjo J, Mellström B, Achaval M, Sassone-Corsit P (1991) Molecular pathways of pain: Fos/Jun-mediated activation of a noncanonical AP-1 site in the prodynorphin gene. Neuron 6:607–617.

Nguyen MQ, von Buchholtz LJ, Reker AN, Ryba NJP, Davidson S (2021) Single nucleus transcriptomic analysis of human dorsal root ganglion neurons. bioRxiv:2021.2007.2002.450845.

North RY, Li Y, Ray P, Rhines LD, Tatsui CE, Rao G, Johansson CA, Zhang H, Kim YH, Zhang B, Dussor G, Kim TH, Price TJ, Dougherty PM (2019) Electrophysiological and transcriptomic correlates of neuropathic pain in human dorsal root ganglion neurons. Brain 142:1215–1226.

Odem MA, Bavencoffe AG, Cassidy RM, Lopez ER, Tian J, Dessauer CW, Walters ET (2018) Isolated nociceptors reveal multiple specializations for generating irregular ongoing activity associated with ongoing pain. Pain 159:2347–2362.

Pertea M, Pertea GM, Antonescu CM, Chang T-C, Mendell JT, Salzberg SL (2015) StringTie enables improved reconstruction of a transcriptome from RNA-seq reads. Nature biotechnology 33:290–295.

Pitcher MH, Von Korff M, Bushnell MC, Porter L (2018) Prevalence and Profile of High-Impact Chronic Pain in the United States. J Pain.

Price TJ, Gold MS (2018) From Mechanism to Cure: Renewing the Goal to Eliminate the Disease of Pain. Pain medicine 19:1525–1549.

Price TJ, Ray PR (2019) Recent advances toward understanding the mysteries of the acute to chronic pain transition. Curr Opin Physiol 11:42–50.

Price TJ, Basbaum AI, Bresnahan J, Chambers JF, De Koninck Y, Edwards RR, Ji R-R, Katz J, Kavelaars A, Levine JD (2018) Transition to chronic pain: opportunities for novel therapeutics. Nature Reviews Neuroscience 19:383–384.

Ray P, Torck A, Quigley L, Wangzhou A, Neiman M, Rao C, Lam T, Kim JY, Kim TH, Zhang MQ, Dussor G, Price TJ (2018) Comparative transcriptome profiling of the human and mouse dorsal root ganglia: an RNA-seq-based resource for pain and sensory neuroscience research. Pain 159:1325–1345.

Ray PR, Khan J, Wangzhou A, Tavares-Ferreira D, Akopian AN, Dussor G, Price TJ (2019) Transcriptome Analysis of the Human Tibial Nerve Identifies Sexually Dimorphic Expression of Genes Involved in Pain, Inflammation, and Neuro-Immunity. Front Mol Neurosci 12:37.

Renthal W et al. (2021) Human cells and networks of pain: Transforming pain target identification and therapeutic development. Neuron 109:1426–1429.

Rigaud M, Gemes G, Barabas M-E, Chernoff DI, Abram SE, Stucky CL, Hogan QH (2008) Species and strain differences in rodent sciatic nerve anatomy: implications for studies of neuropathic pain. Pain 136:188–201.

Rosen S, Ham B, Mogil JS (2017) Sex differences in neuroimmunity and pain. J Neurosci Res 95:500–508.

Rostock C, Schrenk-Siemens K, Pohle J, Siemens J (2018) Human vs. Mouse Nociceptors - Similarities and Differences. Neuroscience 387:13–27.

Rudjito R, Agalave NM, Farinotti AB, Lundback P, Szabo-Pardi TA, Price TJ, Harris HE, Burton MD, Svensson CI (2021) Sex- and cell-dependent contribution of peripheral high mobility group box 1 and TLR4 in arthritis-induced pain. Pain 162:459–470.

Schindler C, Levy DE, Decker T (2007) JAK-STAT signaling: from interferons to cytokines. Journal of Biological Chemistry 282:20059–20063.

Schwab JM, Frei E, Klusman I, Schnell L, Schwab ME, Schluesener HJ (2001) AIF-1 expression defines a proliferating and alert microglial/macrophage phenotype following spinal cord injury in rats. Journal of neuroimmunology 119:214–222.

Serra J, Collado A, Sola R, Antonelli F, Torres X, Salgueiro M, Quiles C, Bostock H (2014) Hyperexcitable C nociceptors in fibromyalgia. Ann Neurol 75:196–208.

Shiers S, Klein RM, Price TJ (2020) Quantitative differences in neuronal subpopulations between mouse and human dorsal root ganglia demonstrated with RNAscope in situ hybridization. Pain 161:2410–2424.

Shiers SI, Sankaranarayanan I, Jeevakumar V, Cervantes A, Reese JC, Price TJ (2021) Convergence of peptidergic and non-peptidergic protein markers in the human dorsal root ganglion and spinal dorsal horn. J Comp Neurol 529:2771–2788.

Singh RP, Hahn BH (2020) Interferon Genes are Influenced by Sex Hormones (17b-estradiol) in SLE. In: Am Assoc Immnol.

Sommer C, Leinders M, Uceyler N (2018) Inflammation in the pathophysiology of neuropathic pain. Pain 159:595–602.

Sorge RE et al. (2015) Different immune cells mediate mechanical pain hypersensitivity in male and female mice. Nat Neurosci 18:1081–1083.

Taga T (1996) Gp130, a shared signal transducing receptor component for hematopoietic and neuropoietic cytokines. J Neurochem 67:1–10.

Tavares-Ferreira D, Shiers S, Ray PR, Wangzhou A, Jeevakumar V, Sankaranarayanan I, Cervantes A, Reese JC, Chamessian A, Copits BA, Dougherty PM, Gereau RW, Burton MD, Dussor G, Price TJ (2021) Spatial transcriptomics reveals unique molecular fingerprints of human nociceptors. bioRxiv:2021.2002.2006.430065.

Vaso A, Adahan HM, Gjika A, Zahaj S, Zhurda T, Vyshka G, Devor M (2014) Peripheral nervous system origin of phantom limb pain. Pain 155:1384–1391.

Wangzhou A, Paige C, Neerukonda SV, Naik DK, Kume M, David ET, Dussor G, Ray PR, Price TJ (2021) A ligand-receptor interactome platform for discovery of pain mechanisms and therapeutic targets. Sci Signal 14.

Ye S, Pang H, Gu Y-Y, Hua J, Chen X-G, Bao C-D, Wang Y, Zhang W, Qian J, Tsao B (2003) Protein interaction for an interferon-inducible systemic lupus associated gene, IFIT1. Rheumatology 42:1155–1163.

Yousuf MS, Price TJ (2020) The importins of pain. Science 369:774–775.

Yu X, Liu H, Hamel KA, Morvan MG, Yu S, Leff J, Guan Z, Braz JM, Basbaum AI (2020) Dorsal root ganglion macrophages contribute to both the initiation and persistence of neuropathic pain. Nat Commun 11:264.

