## Supplemental Figures for "RNA Profiling of Neuropathic Pain-Associated Human DRGs Reveal Sex-differences in Neuro-immune Interactions Promoting Pain"

**Supplemental material**

**Supplementary figures**

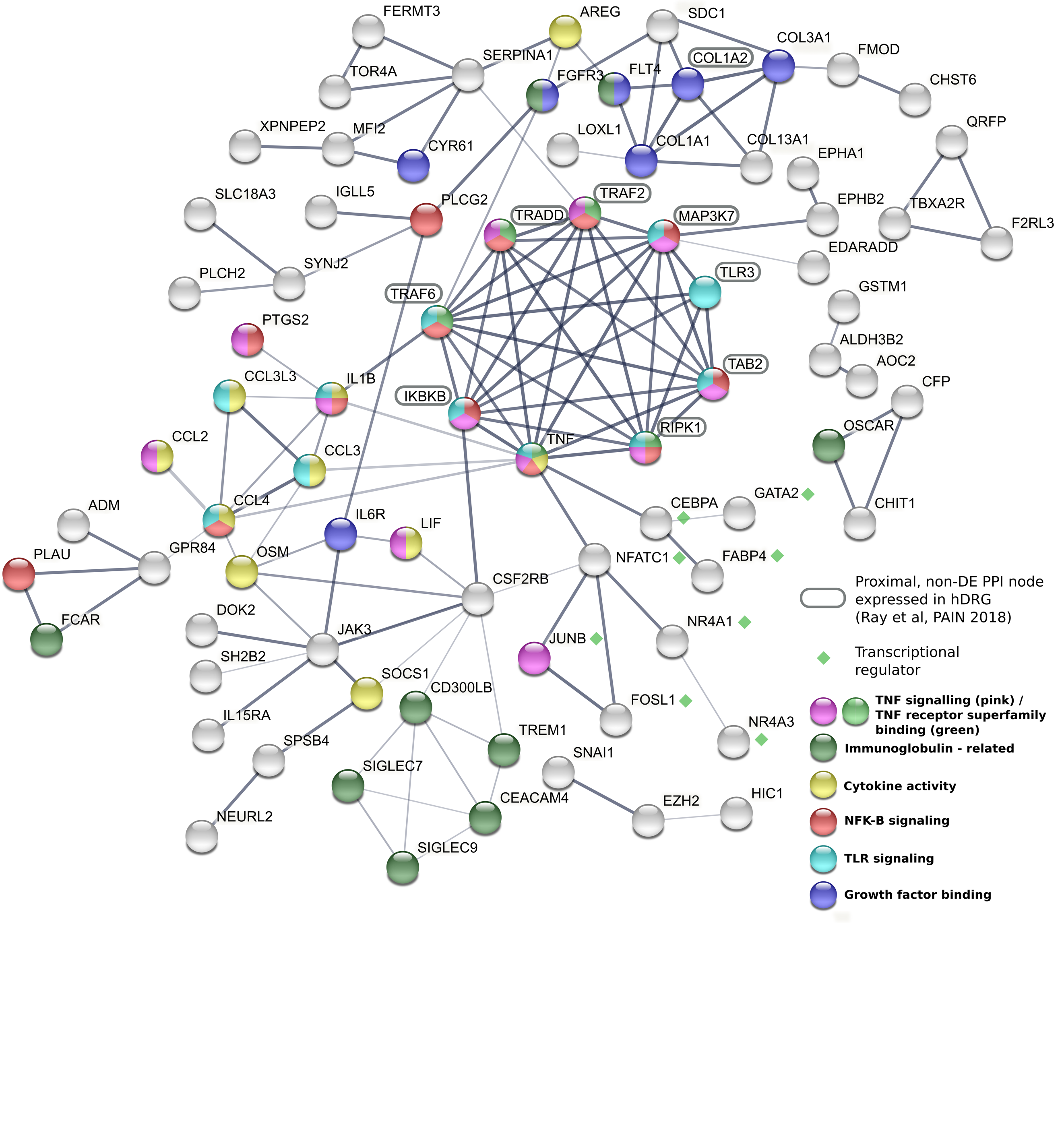

**Supplementary Figure S1.** **StringDB-based protein interaction network for male samples.** Putative protein interaction network based on the StringDB database, seeded with gene products of male pain-associated genes, *OSM* co-expression module and hDRG-enriched genes show distinct evidence of pro-inflammatory cytokine signaling (IL1B, CCL3, CCL4, LIF, SOCS1) and TNF signaling (TNF, IL1B, JUNB, CCL2).

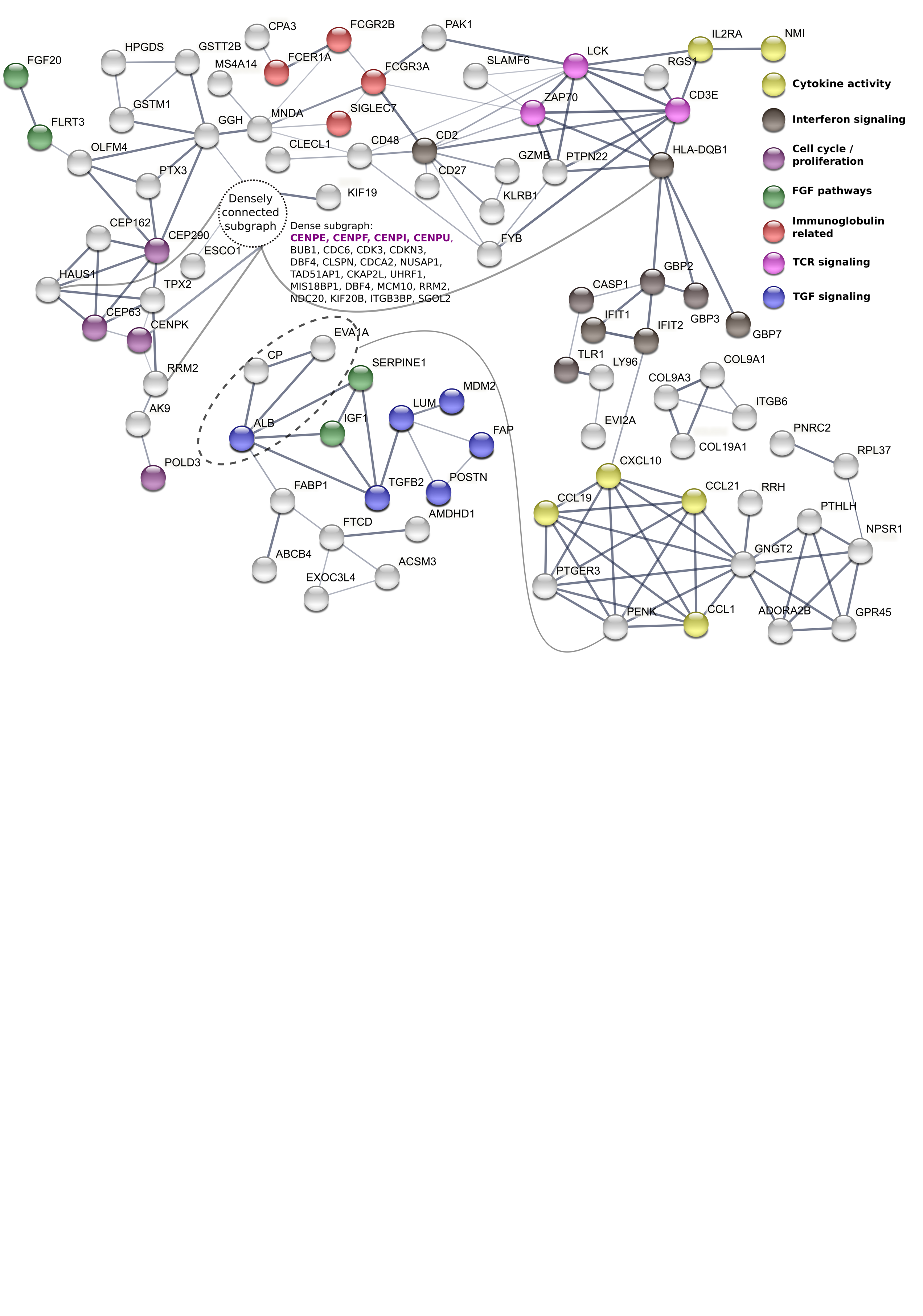
**Supplementary Figure S2.** **StringDB-based protein interaction network for female samples.** Putative protein interaction network based on the STRINGdb database, seeded with gene products of female pain-associated genes, *IFIT1* co-expression module and hDRG-enriched genes show distinct evidence of cytokine signaling (CCL1, CCL19, CCL21), interferon signaling (IFIT1, IFIT2, CASP1, TLR1), and centrosomal proteins (CENPK, CEP63, POLD3).

**Supplemental files**

**Supplemental File 1.** Full patient information table.

**Supplemental File 2.** Sheets A – E for whole transcriptome gene abundance quantification, and sample quality control

**Supplemental File 3.** Sheets A – J for pain-associated gene lists, and membership lists for co-expression modules of *OSM* and *IFIT1.*
